# High-throughput genomic feature extraction reveals environmental adaptations of prokaryotes

**DOI:** 10.64898/2026.09.01.748597

**Authors:** Maria B. Walter Costa, Rose Brouns, Maria Schreiber, Aristeidis Litos, Francesco Bisiach, Heyde Francielle do Carmo França, Casey R.J. Hubert, Manja Marz, Bas E. Dutilh

## Abstract

Understanding the adaptations of microorganisms to their environment is key to predicting the stability and dynamics of microbial communities. To uncover molecular mechanisms of environmental response, we extracted genomic features from 13,554 prokaryotic isolates, and trained machine learning models to identify which ones are most strongly associated with the microbial salinity, temperature, oxygen, and pH preferences. To extract these features in high throughput, including gene families, non-coding RNAs (ncRNAs), oligonucleotides, and amino acid usage, we built FxTractor, a scalable and adjustable pipeline available at: https://github.com/MGXlab/FxTractor. We validated the performance of our models with experimental data from a newly isolated deep-sea extremophile belonging to the genus *Limnochorda* that is not well-represented among the ML training sets, showing strong agreement between predictions and the conditions used to isolate this strain. Our analysis revealed specific gene and ncRNA families associated with each of the four environmental parameters, uncovering both established and potentially new molecular mechanisms. Examples include the bacterial large Signaling Recognition Particle in isolates that are able to grow at high temperatures (≥55°C), suggesting a role in translational pausing and structural stability under thermal stress. We also found the anti-hemB ncRNA to be associated with low-salinity (<0.7% NaCl), indicating a conserved antisense mechanism regulating the energetic costs of heme biosynthesis. Together, these findings provide new insights into microbe-environment interactions, and show how FxTractor enables high throughput discovery of genomic associations.

**Importance:** Microbes have different growth preferences and understanding how this is reflected in the genome is central to microbial ecology and biotechnology. Genomic signatures of environmental adaptation, such as salinity, temperature, oxygen, and pH, may vary from specific encoded proteins and non-coding RNAs to nucleotide and amino acid usage. To analyse these, we developed FxTractor, a flexible open-source bioinformatics pipeline to extract genomic features, that may be readily extended with additional feature mining tools in the future. After applying it to >13,000 prokaryotic genomes, we used machine learning to identify which features were associated with growth preferences, recovering both known and previously uncharacterized mechanisms, including little-researched non-coding RNAs. This work provides both a reusable, extendable tool for genomic feature mining, and new biological insights into the molecular basis of environmental adaptation.

## Introduction

Microbial growth is constrained by the physical environment, where factors such as salinity, temperature, oxygen availability, and pH set the boundary conditions for life. Genomically encoded mechanisms affect where microbes can persist, and how they interact with their environment (Li 2025, Martiny 2015, Philippot 2024). In a context of climate change, with increasing fluctuations of temperature and other environmental factors, identifying microbial preferences and the traits that enable microbial activity under different conditions is critical to predicting ecosystem stability (Ramoneda 2024, Chen 2023, Litalien 2020). Underlying molecular mechanisms may also be relevant for application in industrial and biotechnological contexts (Collette 2025). To define growth preferences, cultivation-based assays remain the benchmark, but these efforts are slow, costly, and have limited throughput (Pham 2012). With the rapid expansion of sequencing data, high-throughput genomics and metagenomics now make it possible to study hundreds to thousands of organisms simultaneously (Weimann 2016). Although progress is hampered by the scarcity of phenotype metadata, databases such as BacDive and PhageDive are closing this gap with experimentally validated cultivation information (Schober 2025, Rolland 2025). In this context, machine learning (ML) provides a powerful framework to link genomic features with phenotypic traits, enabling large-scale prediction of microbial growth requirements beyond the limits of experimental approaches (Dutilh 2013).

High-quality metadata combined with advanced ML techniques, has facilitated capturing genotype-phenotype associations (Dutilh 2013, Goodswen 2021, Alneberg 2020), and predicting growth preferences for environmental parameters such as salinity (Wu 2024, Barberán 2017), temperature (Collette 2025, Li 2019, Sauer 2019), oxygen (Jabłońska 2019, Weimann 2016), and pH (Gado 2025, Ramoneda 2023). ML models may be used to predict laboratory cultivation conditions (Koblitz 2025, Barnum 2024, Oduwole 2025, Liu 2025), and improve our understanding of environmental niches (Alneberg 2020). Recent studies have focused on extracting genomic features including protein coding genes, amino acid frequencies, protein families (Pfam), eggNOG Clusters of Orthologous Groups (COGs), and KEGG annotations (Koblitz 2025, Gado 2025, Barnum 2024, Barberán 2017, Wu 2024).

Interactions of microbes with their surrounding environment involve complex traits. Halophiles, organisms living in high salinity, are efficient when balancing osmosis. They have strategies such as “salt-in”, which are based on production or transportation of compatible solutes (Wu 2024). As an alternative, halophiles can also make use of osmoprotectants, such as sugars, betaines, or amino acids (Lach 2021, Edbeib 2016). Low or high temperatures also select for adaptations, including increasing structural stability of proteins via chaperones. This can lead to interesting industrial applications, such as the production of enzymes with high catalytic capabilities that are resistant to extreme conditions (Dumorné 2017). Oxygen imposes different stresses to cells, such as reactive oxygen species (ROS), which are intermediates of oxygen reduction and can cause damage to DNA and proteins. To mitigate this risk, prokaryotes use ROS scavenging and defense systems (Johnson and Hug 2019). Many of these adaptations depend on protein coding genes, such as ion transporters and chaperones, which can be detected as genomic features, making them relevant for predictive modeling. Microbes in extreme pH conditions deal with acid, oxidative, osmotic, and heavy metal stresses. To maintain homeostasis, they establish transmembrane ion gradients, maintain cell integrity and DNA stabilization, and synthesize osmoprotectants (Aliyu 2024).

While most research has focused on protein-coding genes, non-coding RNAs (ncRNAs) are also important mediators of microbial adaptations (Kohli 2025). These molecules, ranging from small RNAs to complex ribozymes, regulate gene expression at both transcriptional and translational levels and can enable faster responses to environmental fluctuations than protein-mediated signaling pathways (Shimoni 2007). For example, RNA thermometers are ncRNAs that sense temperature changes and regulate transcription and translation to maintain cellular homeostasis under temperature-associated stress (Abduljalil 2018, Roßmanith and Narberhaus 2016). Sodium riboswitches can sense increased sodium concentrations and regulate the expression of genes involved in osmotic stress mitigation. These RNAs also employ mechanisms that exploit sodium accumulation for ATP production (White 2022). Growing evidence highlights the active role of small ncRNAs in oxygen stress responses (Solar Venero 2022, Tan 2023, Dutta and Srivastava 2018). Despite their functional importance, ncRNAs remain underexplored in predictive models.

In this study, we explored ML models to better understand the association of genomic features of prokaryotes with four key environmental parameters: salinity, temperature, oxygen, and pH. We developed a high-throughput feature extraction pipeline, FxTractor, and applied it to a dataset of 13,554 isolates with experimentally determined growth preferences for these four parameters. The versatility of FxTractor associated with ML models allowed us to compare four different genomic feature types, not only benchmarking their predictive performance, but also to identify specific mechanisms of adaptation, underpinning responses to environmental factors. Our work brings new insights to better understand how microbes interact with the environment.

## Material and Methods

All tools were run with default parameter settings unless mentioned otherwise. Scripts were developed with Python 3.13.3 and Jupyter Notebook 7.4.2. Figures were made with matplotlib, seaborn, and draw.io, and visualization was improved with Inkscape. The repository https://github.com/MGXlab/abiotic_environment contains scripts, a file with information about the analysed isolates and metadata details, and a CSV file with the results of the Shapley additive explanations (SHAP) analysis.

### Datasets and curation

To obtain information about growth preferences of prokaryotes, we downloaded metadata for 91,228 microbial isolates (90,241 bacteria, 987 archaea, **Supplementary Table 1**) from BacDive (Schober 2025) and extracted information about the salinity, temperature, oxygen, and pH preferences. We removed all entries that had ambiguous or unclear descriptions, and only kept entries that clearly indicated growth in the lab (data fields “growth” or “tested relation” matching either “positive growth" or “positive optimum”). In the case of oxygen, we only kept entries showing clear aerobe or anaerobe profiles (data field “oxygen tolerance” matching “aerobe”, “obligate aerobe”, “anaerobe” or “obligate anaerobe”).

Next, we prepared the data for Machine Learning (ML) regression and classification modeling. Since isolates could contain more than one reported value of a given environmental factor, we considered the mean value when building regression models. In the case of classification models, we used contrasting classes. For salinity, the low class had mean NaCl ≤0.7% and the high class ≥3.5%. For temperature, the low class had mean temperature ≤17°C and the high class ≥55°C. To build balanced classes, we randomly undersampled the class with the highest number of samples so that it matched the number of samples of the class with the lowest number of samples. In the case of pH, we chose not to build classes but only worked with regression models, since most isolates had pH values between seven and eight (**Figure 1**), and the data did not neatly fall into two classes.

**Figure 1.**
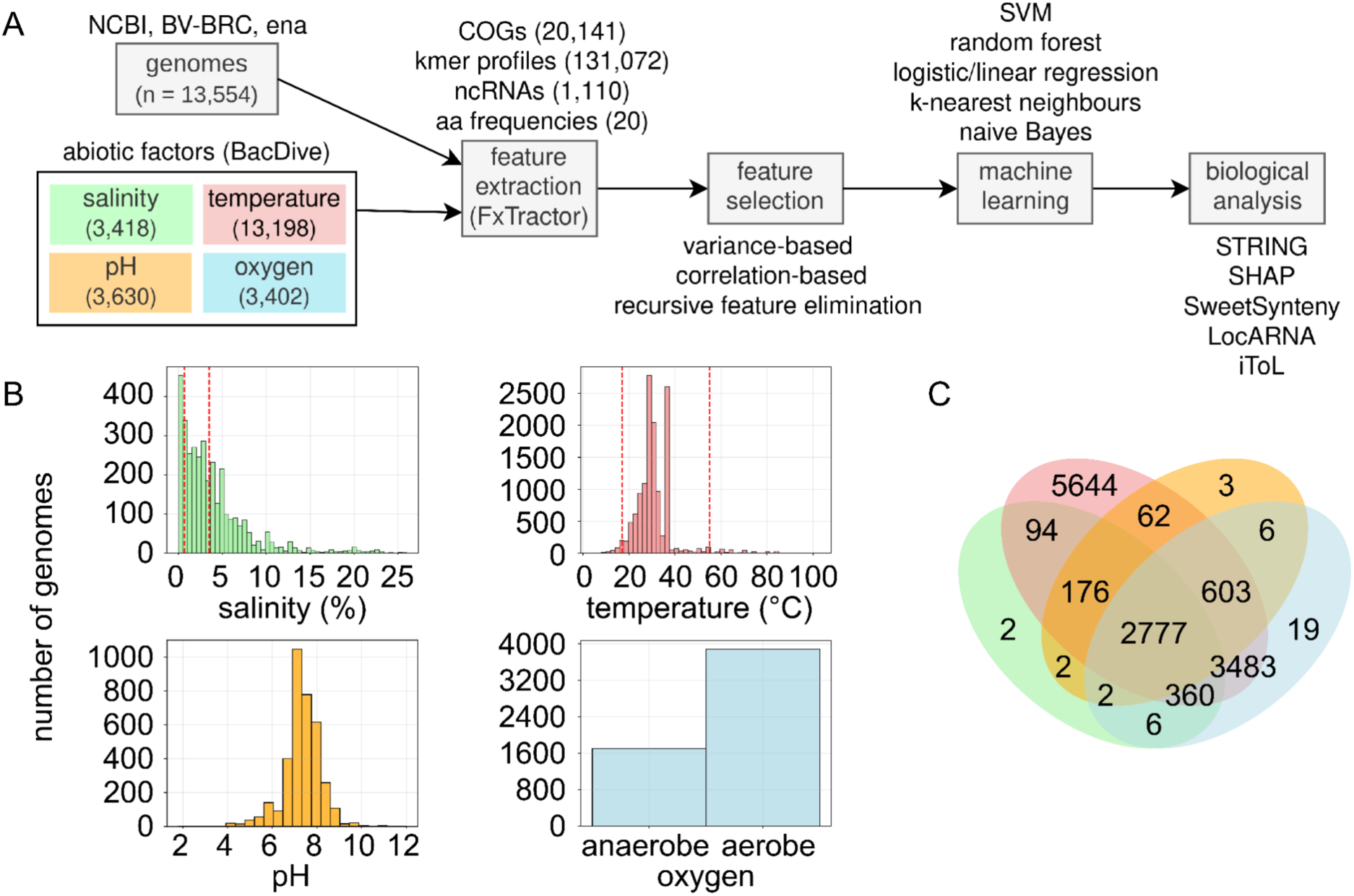
Study pipeline and dataset distribution. **(A)** Pipeline for linking genomic features to environmental parameters. In total, we processed n=13,554 prokaryotic genome sequences with high-quality metadata annotations of salinity temperature, oxygen, and/or pH. Then, we extracted genomic features with the FxTractor pipeline and performed biological analysis on features considered important by ML models. **(B)** Number of genomes analyzed for different environmental parameters. Red dashed vertical lines show thresholds for creating low and high classes. **(C)** Summary of isolates with overlapping metadata.

To ensure high-quality genomic data, we downloaded all 38,822 genome sequences marked as “complete”. Note that the "complete" genomes still contained between 1- 2,971 contigs (mean: 54 ± 115). As BacDive reported assemblies in different databases: NCBI Genome database (Sayers 2022), BV-BRC (Olson 2023), and IMG (Chen 2021), we downloaded sequences from these three repositories (n=16,879, 15,364, and 6,579, respectively). Next, we filtered out 25,268 genomes that did not have metadata on salinity, temperature, oxygen, pH or and those with CheckM (Parks 2015) completeness ≤90% or contamination >5%. For isolates with multiple assemblies, we selected the one with the highest completeness. If two assemblies had the same completeness, we chose the one with the fewest contigs. This yielded a dataset of 13,554 genomes (available at https://github.com/MGXlab/abiotic_environment).

### FxTractor feature extraction pipeline

To obtain genomic features from the genome sequences, we extracted: eggNOG Clusters of Orthologous Groups (COGs) (Cantalapiedra 2021), nucleotide usage (9 nt kmers), Rfam ncRNA families, and amino acid frequencies (**Table 1**). COGs and ncRNA families were represented in each genome as absent or present (0 or 1), and kmers as the absolute number of occurrences. We chose the length of 9 for kmers, since it provides more information than shorter ones and the final matrix of 131,072 kmers was still feasible to process computationally.

**Table 1.** Number of features selected per environmental factor and feature type along with the best performing machine learning model. The number of isolates in classification datasets combines the two contrasting classes (low and high or aerobic and anaerobic). Numbers of features for regression models are described in **Supplementary Table 3**.

| Environmental factor | Feature type | ML model | Isolates with annotation | Initial features | Selected features |
| --- | --- | --- | --- | --- | --- |
| Salinity | COGs | Classification (Logistic Regression) | 932 | 20,141 | 3,002 |
|  | kmers | Classification (SVM, Linear Kernel) | 932 | 131,072 | 1,873 |
|  | ncRNAs | Classification (SVM, Linear Kernel) | 932 | 1,110 | 189 |
|  | Amino acids | Classification (SVM, Polynomial Kernel) | 932 | 20 | 20 |
| Temperature | COGs | Classification (SVM, Linear Kernel) | 668 | 20,141 | 3,173 |
|  | kmers | Classification (SVM, Linear Kernel) | 668 | 131,072 | 9,062 |
|  | ncRNAs | Classification (SVM, Linear Kernel) | 668 | 1,110 | 149 |
|  | Amino acids | Classification (SVM, Polynomial Kernel) | 668 | 20 | 20 |
| Oxygen | COGs | Classification (SVM, Linear Kernel) | 3,402 | 20,141 | 4,823 |
|  | kmers | Classification (SVM, Linear Kernel) | 3,402 | 131,072 | 8,770 |
|  | ncRNAs | Classification (SVM, Linear Kernel) | 3,402 | 1,110 | 961 |
|  | Amino acids | Classification (Random Forest) | 3,402 | 20 | 20 |
| pH | COGs | Regression<br>(SVM, Polynomial<br>Kernel) | 3,630 | 20,141 | 3,638 |
|  | kmers | Regression<br>(Random Forest) | 3,630 | 131,072 | 4,096 |
|  | ncRNAs | Regression<br>(knn regressor) | 3,630 | 1,110 | 737 |
|  | Amino<br>acids | Regression<br>(Random Forest) | 3,630 | 20 | 20 |

Features were extracted with the FxTractor pipeline, which we developed using the Snakemake platform (Mölder 2021). The pipeline uses as default 60 logical cores for Jellyfish 1.1.12 (Marçais and Kingsford 2011), CheckM 1.2.2 (Kang 2019), eggNOG emapper 2.1.11 (Cantalapiedra 2021), and cmscan of Infernal 1.1.5 (Nawrocki and Eddy 2013). All other rules use 2 logical cores. All tools run with their default parameters within the pipeline, with the exception of eggNOG emapper, for which an e-value filter was included of 0.0001. To decrease the run time of emapper, we set its block size to 10.0. To improve predictions of non-coding RNAs (ncRNAs), we included a filter that only kept cmscan predictions of prokaryotic families (based on the Rfam lists: (https://github.com/Rfam/rfam-taxonomy/blob/master/domains/archaea.csv and https://github.com/Rfam/rfam-taxonomy/blob/master/domains/bacteria.csv). We also removed hits with e-value >0.001 and bias>50.

For benchmarking requirements of run time and memory, we ran the pipeline with the settings mentioned above for 100 random genomes. The main Snakefile and all individual rules, such as for eggNOG emapper or cmscan of Infernal, ran in dual-socket Intel Xeon Gold 8360Y CPUs (96 logical cores, 24 cores per socket, two threads per core), 256 GB RAM, on a Linux operating system (kernel version 4.18.0-425.13.1.el8_7.x86_64). During execution, up to 96 CPUs and 188 GB of memory per node could be allocated. When the pipeline was first run on our cluster, the initial installation of all Conda environments took 2h 45m. This installation is required only once. For subsequent runs, the pipeline can proceed directly to feature extraction without repeating the setup.

### Feature selection

To reduce the feature space and select the most important features, we applied feature selection to COGs, ncRNAs, and kmers. Feature selection consisted of three steps: (i) removing features with low variance, (ii) joining highly correlating features, and (iii) applying recursive feature elimination (RFE). To choose an appropriate threshold for filtering out features with low variance, we performed a benchmark based on ten iterations of 5-fold cross-validation Random Forest (RF). The threshold with the highest median cross-validation metric was chosen per environmental factor and feature type (COGs, kmers, and ncRNA families, **Supplementary Table 2** and **Supplementary Figure 1**). Metrics were: F1 score for classification and maximum absolute error (MAE) for regression. Variance was measured for each feature type using the numpy function var, which measures the spread of a distribution. Afterwards, variances were scaled to a range [0; 1] to make different features comparable.

To reduce the number of highly correlating features that could confound ML-based feature scoring, we clustered them using single linkage based on Spearman correlation and chose the feature with the highest mean correlation value to the other group members to represent the group. To find appropriate correlation thresholds for each dataset (**Supplementary Table 2** and **Supplementary Figure 2**), we used a similar benchmark strategy as for variance-based selection (above).

The third selection step was performed using RFE based on decision trees. For that, we calculated feature importance scores and plotted them in a histogram (**Supplementary Figure 3**). We then defined “elbow points” as the point in the histogram where the scores exhibited a sharp increase of ∼90%. For datasets showing clear “elbow-points”, we kept all features with scores above the one at the “elbow point” (**Supplementary Table 2**). For datasets not showing an “elbow point”, we kept all features. Lasso, L2, and tree-based methods were also applied to the datasets, but only RFE showed “elbow-points”, facilitating the choice of how many features to remove.

After performing all three feature selection steps, we defined as final datasets (**Supplementary Table 3**) those yielding the best RF performance in 5-fold cross-validation analysis (**Supplementary Figure 4**). For the kmer regression datasets, which contained a substantially larger number of features (**Supplementary Table 3**), we selected the step that removed the greatest number of features while maintaining very similar performance (**Supplementary Figure 4**). This approach ensured computational feasibility without compromising model quality.

### Machine Learning modelling

Next, we built ML regression and classification models to identify to what extent the genomic features extracted and selected above were associated with salinity, temperature, oxygen, and pH preferences, as annotated for the BacDive isolates. The aim was to predict the growth values (regression) or category (high or low, classification) of salinity, temperature, and pH, calculated as the mean of all corresponding BacDive records.

To identify the best-performing ML methods with our selected features, we first carried out a preliminary comparison of linear and non-linear models using 5-fold cross-validation, including linear/logistic regression, RF, k-nearest neighbours, and Support Vector Machine (SVM) with linear, polynomial, and Radial Basis Function (RBF) kernels. This yielded best-performing models for each of the datasets, considering the four environmental parameters, four feature types (kmers, COGs, ncRNAs, and amino acid frequencies), and ML problem type (classification or regression). For oxygen there is only a classification model (no regression) due to its categorical nature, whereas for pH there is only a regression model (no classification) due to the mostly neutral pH values and difficulty creating contrasting classes. To contextualize the model performance, we also modeled trivial predictors using the Dummy functions of sci-kit learn as follows. For regression models, we include the median dummy function, which predicts for every case the median value of the training set. For classification models, we include a random class with equal probability of assignment as a dummy function.

After selecting the best ML method from the preliminary step above, we then optimized the corresponding model for each dataset by tuning the hyperparameters: logistic regression (’C’: np.linspace(0.1, 1.0, 10)), RF (’n_estimators’: [500], ‘max_features’:

[10, 15, 20], ‘min_samples_leaf’: [5, 8,10]), knn regressor (’n_neighbors’: [3, 5, 7, 9, 11, 13, 15]), and SVM (linear or polynomial kernel, ‘C’: np.linspace(0.1, 1.0, 10)). Optimal models per dataset can be found in **Supplementary Table 4**.

We assessed the performance of the regression models using R^2^ values and of the classification models using the F1-score. For the best models, we used SHAP (Lundberg 2018) to rank features by importance.

### Validation of models with an environmental isolate

To test our prediction models on a novel isolate with a genome that was not included in the training set, we isolated and sequenced the genome of an endospore-forming marine extremophile. This bacterial strain was obtained from sediments of a natural oil seep in the abyssal plain of Eastern Gulf of Mexico (27.91°N, 86.75°W; 2,929 m water depth). Enrichment cultures were established in triplicate by anoxically mixing autoclaved marine sediment (three consecutive cycles of 121 °C for 20 min) with sterile artificial seawater medium (pH 7.3, NaCl 3.5%), supplemented with 20 mM sulfate and a 5 mM mixture of six short-chain fatty acids (acetate, formate, lactate, propionate, butyrate, and succinate). These sediment slurries were incubated at 50 °C in the dark and subsampled on days 0, 14, 21 and 28 for metabolite analysis and DNA sequencing.

Changes in short-chain fatty acid concentrations over time inpost-autoclave 50°C incubations provided an indication of microbial activity. To measure this, slurry aliquots were centrifuged at 21,100 x g for 5 minutes at room temperature followed by filtering the supernatant through a 0.2 µm syringe filter. This was transferred into glass vials and analysed using a Thermo RS3000 high-performance liquid chromatography (HPLC) fitted with an Ultimate 3000 UV detector set at a wavelength of 210 nm. Separation was achieved over an Aminex HPX-87H organic acid column (BioRad, USA) under isocratic conditions (0.05 mM H_2_SO_4_) at 60°C with a run time of 20 min. Concentration data was processed using Chromeleon version 7 software using valley to valley peak integration algorithms comparing to the retention time of known standards.

Genomic DNA was extracted from 300 µL slurry subsamples using the DNeasy PowerLyzer PowerSoil Kit (Qiagen, USA) according to the instructions provided by the manufacturer except for the inclusion of a 10 min incubation at 70°C immediately after the addition of Solution C1 to enhance cell lysis. We included negative controls for each set of DNA extraction using Milli-Q water in place of slurry subsamples. We quantified the concentrations of extracted DNA using the Qubit dsDNA High Sensitivity assay kit on an Invitrogen Qubit 2.0 Fluorometer (Fisher Scientific, Canada). Triplicate PCR used the primer pair SD-Bact-341-bS17/SD-Bact-785-aA21 modified with Illumina MiSeq overhang adapters (Klindworth 2013) to amplify a 444 bp fragment spanning the V3 and V4 hypervariable regions of the 16S rRNA gene. Each PCR reaction was performed in triplicate to minimize bias and consisted of a 30-cycle program. DNA was initially denatured at 95°C for 5 min, which was followed by 30 cycles of denaturing at 95°C for 30 sec, annealing for 60°C for 45 sec, extension at 72°C for 1 min. The reaction concluded with a final extension at 72°C for 5 min. After each PCR run, we validated samples, DNA blanks and PCR blanks by agarose gel electrophoresis. We pooled and purified the PCR products using a NucleoMag NGS Clean-up and Size Select kit (Macherey-Nagel Inc., USA). Indexed amplicons were normalized to an equimolar amount, loaded on a 96-well plate, and sequenced on an Illumina MiSeq benchtop sequencer using Illumina’s v3 600-cycle reagent kit to obtain 300 bp paired-end reads.

We processed and analysed the raw reads using base R version 4.2.2 (R Core Team 2022). We used the DADA2 package version 1.26.0 to perform primer trimming, quality filtering and amplicon sequence variant (ASV) inference were (Callahan 2016). When filtering, we allowed for up to two expected errors for both the forward and reverse reads and trimmed reads when their quality score dropped below 25. To ensure removal of primer dimers and chimeras, we discarded amplicons outside the 440-446 bp range.

After ASV analysis confirmed near-purity of the enriched strain (see Results), we obtained the genome of the enriched bacterium by pooling extracted DNA from 21-day old cultures and submitting it for shotgun metagenomic sequencing to the Center for Health Genomics and Informatics in the Cumming School of Medicine, University of Calgary. DNA was sheared using a Covaris S2 ultrasonicator (Covaris, USA), and fragment libraries prepared using a NEBNext Ultra II DNA Library Prep Kit for Illumina (New England BioLabs, USA). Metagenomic libraries were sequenced on the Illumina NovaSeq platform (Illumina Inc., USA) using an S4 flow cell with Illumina 300 cycle (2 × 150 bp) V1.5 sequencing kit.

The 25,797,141 raw sequences obtained from the single sample library were quality-checked using FastQC version 5.26.2 before being filtered and trimmed using BBDuk (BBTools suite, http://jgi.doe.gov/data-and-tools/bbtools). We assembled the trimmed and filtered reads using MEGAHIT version 1.2.2 using default parameters and with <500 bp contigs removed (Li 2015). Binning was performed using MetaBAT2 version 2.12.1 (Kang 2019). Lineage, genome completeness and contamination were inferred using CheckM version 1.0.11 (Kang 2019) Taxonomy assignment was performed using GTDB-Tk version 1.3.0 (Chaumeil 2019) with reference data R95. Average nucleotide identity values between the assembled genome and its closest cultured relative was computed using the ANI calculator tool (Goris 2007) We then constructed a phylogenetic tree based on the comparison of multiple single copy genes using GToTree v1.6.31 and visualized it with iTOL (Lee 2019, Letunic and Bork 2021). To plot the assembled genome, we scaffolded it over the genome of its closest cultured relative using RagTag v2.1.0 (Alonge 2022) and performed a pairwise nucleotide identity using FastANI v1.3.4 (Jain 2018). These results were then visualized using Proksee v1.6.1 (Grant 2023). We obtained genomic features for this bacterium with our FxTractor pipeline, predicted the organism’s preferences for different parameters, and compared those predictions with the experimental values.

### Biological analysis of high-ranking ncRNAs

To assess the biological significance of ncRNAs prioritized by regression models and SHAP analysis (see “Machine Learning modelling”), we analyzed their genomic context, conservation, and structural topology as follows. Genomic context was assessed with SweetSynteny 1.0 (https://github.com/rnajena/SweetSynteny, Schreiber 2025) using the genomes with temperature and salinity metadata, where relevant ncRNAs were found (SRP: 657, anti-hem: 54), as described in the “Datasets and curation” section. Due to homology between bacterial large and small SRP, the latter annotation was removed for genomes where annotations overlapped. Because the genomes were already annotated, we set the ‘from_gff’ and ‘ignore_overlap’ search option in SweetSynteny, with ‘adjacent_gene_clustering’ to ‘hmmscan,cmscan’ to account for the wide diversity in species. To identify Signaling Recognition Particle (SRP) in the list of annotations in the GFF file, parameters were used ‘substring_search’. Structural sequence alignments were generated using LocARNA 2.0.0 (Will 2007) with the parameters ‘stockholm’, ‘consensus-structure alifold’, and ‘keep-sequence-order’. Alignments were manually curated in Emacs RALEE mode 0.8 (Griffiths-Jones 2005) and can be found on Github (https://github.com/MGXlab/abiotic_environment). Secondary structures were derived from conserved sequence alignments of Rfam 15.1 and visualized with R2DT (Sweeney 2021) and Jalview (Waterhouse 2009). A taxonomic tree showing the occurrence of anti-hemB was constructed as described in the “Environmental isolate” section. The Newick file was generated using ETE 3 (Huerta-Cepas 2016) and subsequently annotated and visualized with iTOL (Letunic and Bork 2021).

## Results

We set out to identify prokaryotic genes and genomic adaptations associated with microbial growth conditions and environmental parameters, specifically salinity, temperature, oxygen availability, and pH. To obtain datasets with growth preferences, we first downloaded the genome sequences and cultivation metadata of 92,150 isolates. After filtering and curation for isolates with complete genome sequences and growth metadata, we used 3,418 isolates for salinity, 13,198 isolates for temperature, 3,630 isolates for pH, and 3,402 isolates for oxygen (**Figure 1**). In total, the examples analysed span 13,554 total isolates (13,107 bacteria and 447 archaea). From these, we extracted four different genomic feature types: nucleotide usage patterns (kmers, k=9), gene families (eggNOG Clusters of Orthologous Groups, COGs), ncRNA families, and amino acid frequencies in encoded proteins, totalling over 150 thousand genomic features (**Figure 1A**). After filtering and feature selection, we used Machine Learning (ML) to associate these features with salinity, temperature, oxygen, and pH, and analysed the biological significance of the identified associations. The number of genomes with annotations for the different environmental parameters is shown in **Figure 1BC**.

In the following sections, we first present the FxTractor pipeline, which we built to extract diverse genomic features. Then, we describe the ML models to predict environmental factors and compare the predictive power of different feature types.

Next, we test our models with an isolated marine extremophile grown under anoxic, high-temperature conditions in the lab. Finally, to identify genomic adaptations to the environment, we investigate the important features found by our ML models, including COGs and ncRNAs.

### FxTractor: a flexible open-source pipeline for high-throughput genomic feature extraction

We developed the pipeline FxTractor to automatically extract features from prokaryotic genome sequences (available at https://github.com/MGXlab/FxTractor). In our case, the extracted features form the input to our ML models that predict environmental factors. The currently implemented feature types include ters, COGs, ncRNAs, and amino acid frequencies, which we expect to be relevant for adaptations to the growth environment, and could reveal underlying biological mechanisms. FxTractor takes nucleotide FASTA files containing individual draft or complete genome sequences as input, which are screened for genomic features through steps described in **Figure 2**.

**Figure 2.**
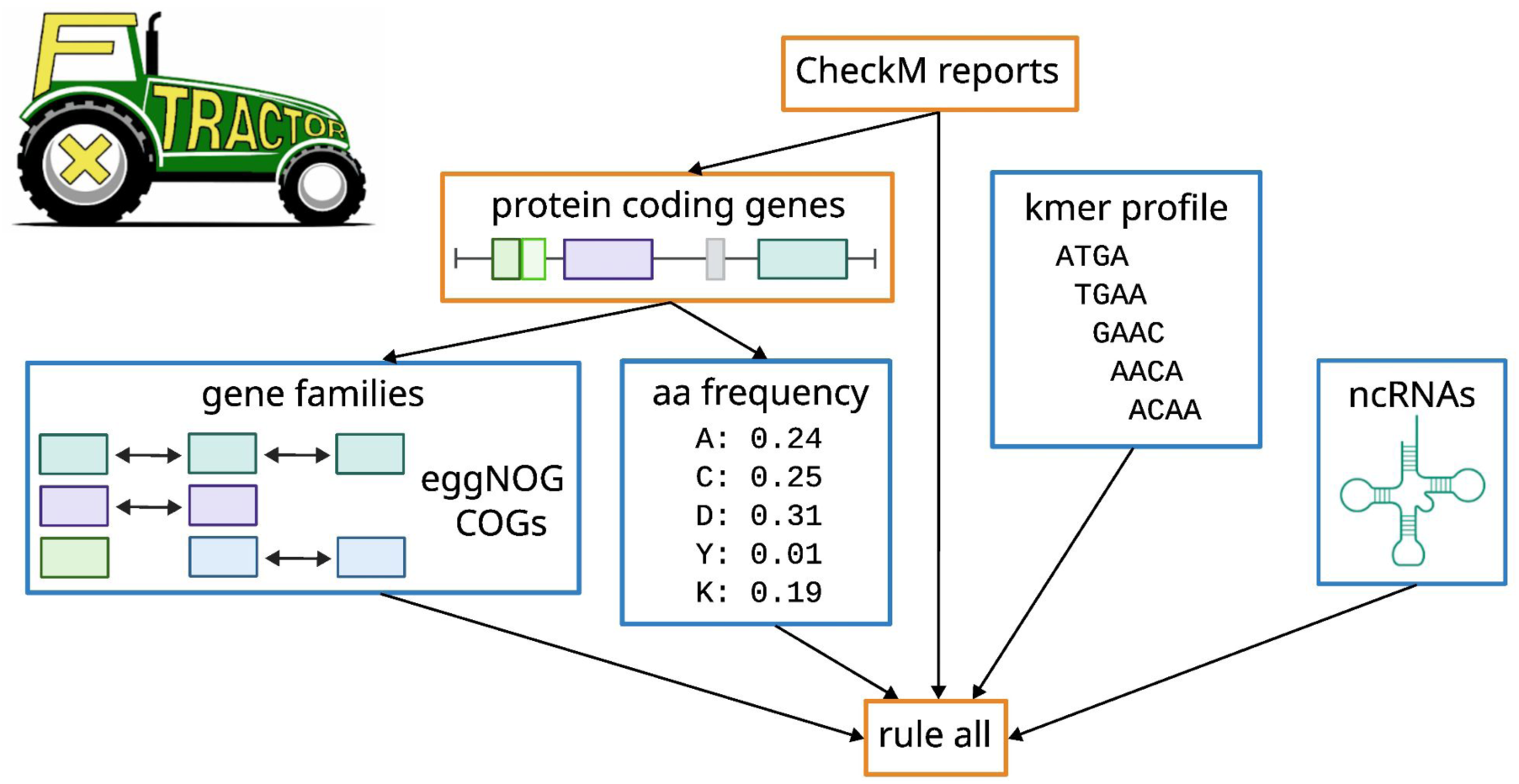
The FxTractor Snakemake pipeline extracts different genomic features from prokaryotic genomes. Internal rules are represented as boxes. Orange boxes represent intermediate rules and the rule “all”, by which the user can define which features to extract. Blue boxes represent features that are produced as output.

As a benchmark for estimating run time and memory requirements, we randomly selected 100 genome sequences from our collection of 13,554 isolates and observed that the required memory usage varied widely between genomic features, from a few megabytes (MB) for kmers to ∼26 gigabytes (GB) for COGs. Larger genomes did not require more memory than smaller ones (**Figure 3A-D**). A similar trend was observed for running time, which ranged from a few seconds to ∼21 minutes, where COGs and ncRNAs, but not kmers took significantly longer to process in larger genomes than in smaller ones (**Figure 3E-H**). Obtaining COGs with eggNOG emapper required the most computational resources (**Figure 3D**). Although emapper implements rapid homology search algorithms, functional annotation of genes is still computationally costly compared to the other features (Grigson 2025, Cantalapiedra 2021).

**Figure 3.**
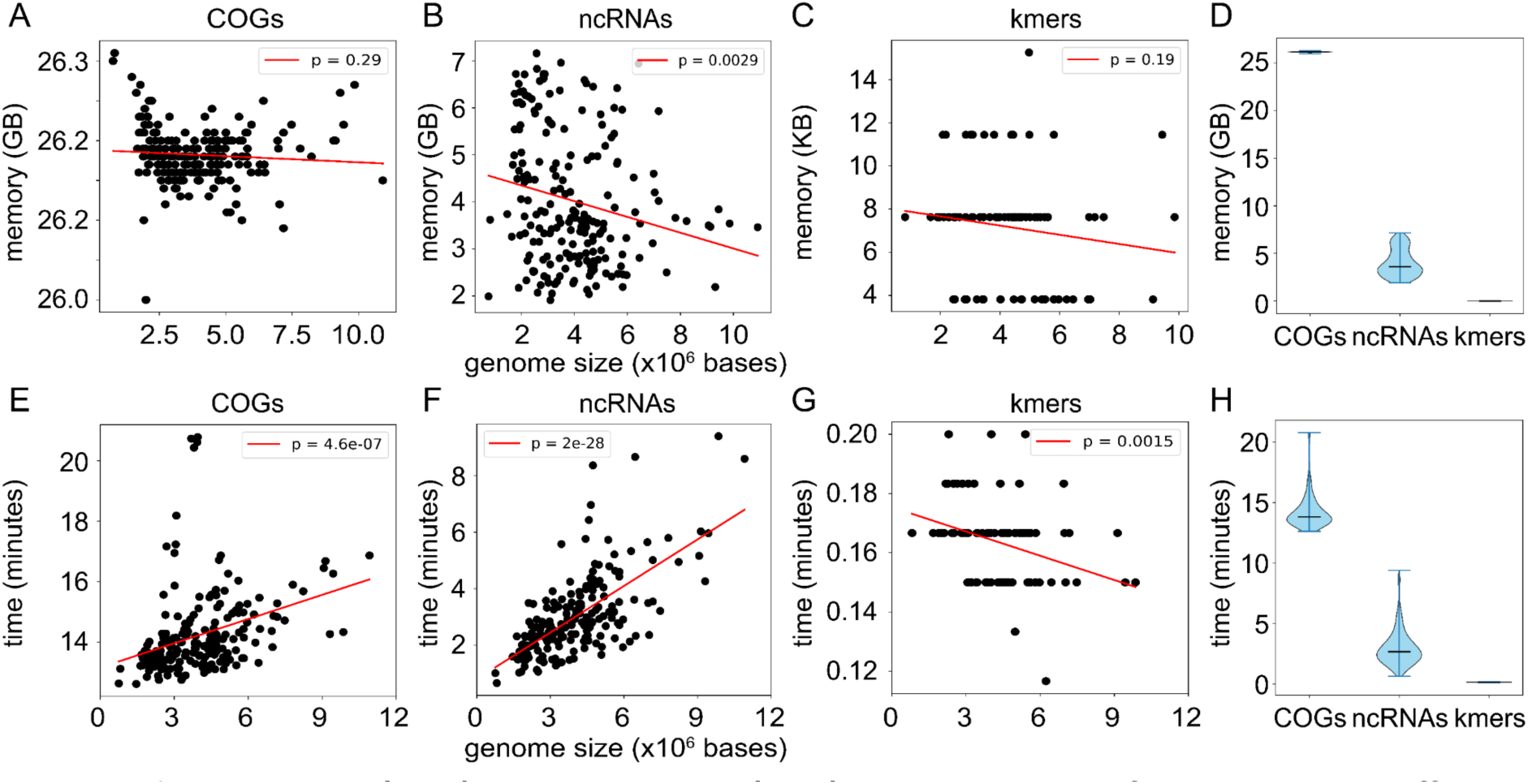
Memory **(A-D)** and runtime **(E-H)** requirements for obtaining different genomic features with FxTractor. Reported values represent calculations on 100 randomly selected prokaryotic genomes between 816,373 and 10,915,467 bases in size (mean: 4,249,162 ± 1,870,084 standard deviation). Red lines represent best-fit linear regressions, with reported p-values indicating whether the regression slope differs from zero. Violin plots in panels **C** and **D** summarize the per-genome memory and runtime requirements, respectively, with median values as black horizontal lines.

The Snakemake implementation facilitates deployment of FxTractor on a high-performance computing cluster (HPC) for automatic parallelization of large numbers of genomes. We tested the pipeline in the Slurm Workload Manager, but Snakemake is also compatible with other HPC systems. Within the pipeline, we specified conda environments using YAML files, which list the required software and their versions. This allows for automatic software installation and full reproducibility.

### Machine learning models to predict environmental factors

With our feature extraction pipeline (**Figure 2**), we obtained 131,072 kmers (k = 9), 20,142 COGs, 1,110 ncRNA families, and 20 amino acid frequencies. Feature selection was an important step to decrease the number of features, while retaining the maximum amount of meaningful information (**Table 1**). Removing low-variance features had the greatest impact, reducing an average of 57.3% of the features (**Supplementary Table 3**).

To obtain optimal ML models for predicting environmental factors, we evaluated seven linear and non-linear algorithms using 5-fold cross-validation and optimized their hyperparameters, selecting the best-performing models for each environmental factor (**Supplementary Table 4**). Our best classification models presented F1-scores of 0.88 for salinity, 0.99 for temperature, and 0.97 for oxygen (**Figure 4A-C**), demonstrating high discrimination between the two contrasting classes. For contextualization, trivial models that assign a random class with equal probability showed F1-scores ∼0.50 (**Figure 4A-C**).

**Figure 4.**
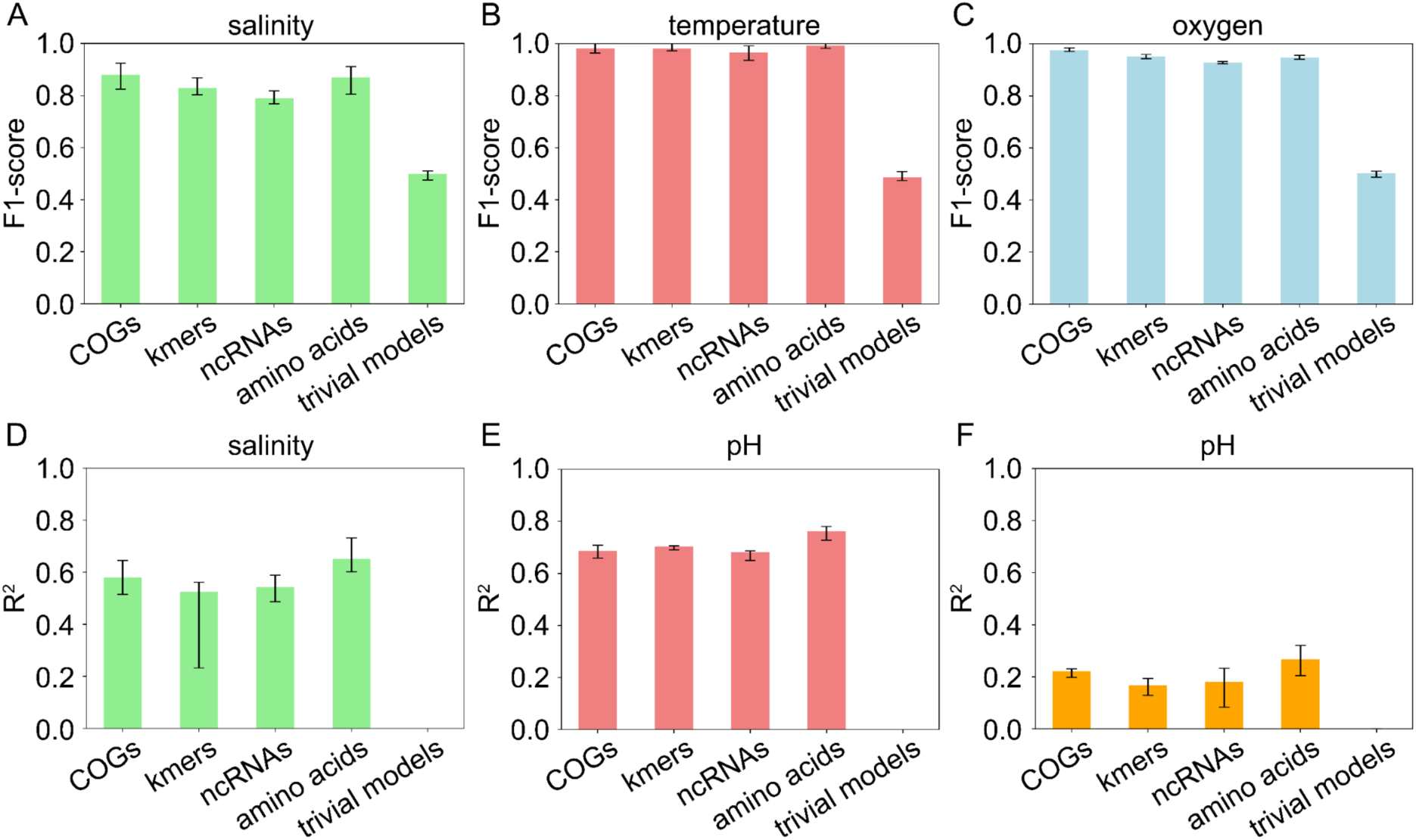
Performance of models for prediction of **(AD)** salinity, **(BE)** temperature, **(C)** oxygen, and **(F)** pH. **(A-C)** classification models, **(D-F)**: regression models. Bars: mean value of five cross-validation slices, whiskers: minimum and maximum values of cross-validation.

Regression models showed differences in predictive power of different feature types, with amino acid frequencies being more predictive than others (**Figure 4D-F**). The R^2^ values of our best models using amino acid frequencies were: 0.65 for salinity, 0.76 for temperature, and 0.27 for pH. Trivial models showed R^2^ values ∼0.0, which are equivalent to predicting values by chance (**Figure 4D-F**). Poorer performance of pH likely reflects that pH values comprise a more uniform dataset (**Figure 1B**), with most isolates growing around pH 7, the remainder being too limited for picking up a meaningful signal. Temperature and salinity values have a wider range, the resulting genomic adaptations providing strong signals for the prediction models. The number of data points also plays a role, with n = 3,418 for salinity, n = 13,198 for temperature, which has the highest performance, and n = 3,630 for pH.

### Models successfully predict environmental parameters for a novel isolate

To validate our regression models, we isolated a thermophilic spore-forming bacterium from a deep-ocean hydrocarbon seep in anoxic artificial seawater medium. The presence of a single highly prevalent bacterial ASV taxonomically assigned to the genus *Limnochorda* was observed in an enrichment culture that was after it was autoclaved and then incubated for 28 days at 50 °C (**Figure 5B**). Lactate and formate, but not sulfate were consumed during the incubation (**Figure 5C**), suggesting that endospores survived autoclaving and then germinated and grew fermentatively. Shotgun metagenomic sequencing on DNA extracted after 21 days incubation resulted in a single bin, with 86.27% genome completeness, 0.98% contamination and 0% strain heterogeneity.

**Figure 5.**
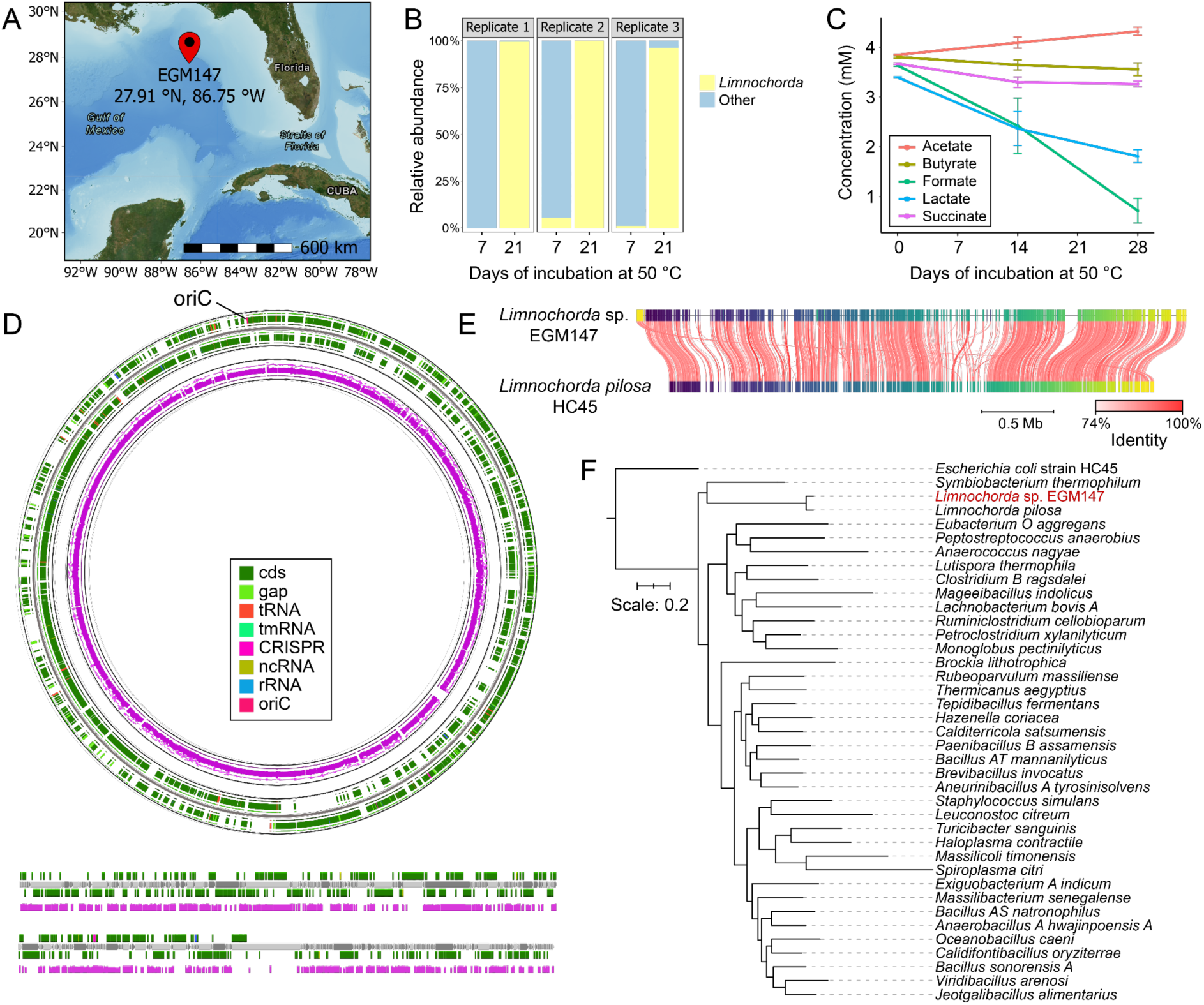
**(A)** Map of the EGM147 sediment sampling site; **(B)** Relative abundance of a single ASV assigned to the Limnochorda genus from the 16S rRNA gene amplicon libraries at day 7 and day 21 of 50 °C anaerobic sediment incubations **(C)** Organic acid concentration in sediment microcosms over 28 days of anaerobic incubation at 50 °C as detected by HPLC measurements. Days of incubation are counted following the three subsequent cycles of autoclaving. Error bars represent the standard deviation of the three replicates; **(D)** Genome map of Limnochorda sp. EGM147 scaffolded against the reference genome of Limnochorda pilosa strain HC45 (ANI = 86.27%). Tracks include scaffolded contigs (magenta) and forward- and reverse-strand CDS (color legend box). Linear contigs that could not be placed during scaffolding are displayed with the same annotation scheme. **(E)** FastANI pairwise nucleotide identity between the scaffolded Limnochorda sp. EGM147 assembly and the L. pilosa HC45 reference genome; color intensity reflects local sequence identity; **(F)** Maximum likelihood phylogenetic tree comparing Limnochorda sp. EGM147 with Limnochorda pilosa, the only cultured representative from the class Limnochordia, cultured representatives from the Bacillota phylum randomly selected from GTDB (Parks 2025), and Escherichia coli strain HC45 (RefSeq: GCF_000008865.2) as an outgroup. The scale bar corresponds to 0.2 amino acid substitutions per alignment position. All bootstrap values were higher than 60%.

Taxonomy assignment against the GTDB-tk database assigned the isolate to the genus *Limnochorda*. There exists only one cultured representative species within the *Limnochordia* class, *Limnochorda pilosa* (Watanabe 2015), which shares 86.05% average nucleotide identity to our isolate (**Figure 5EF**). This new isolate, referred to here as *Limnochorda* sp. EGM147 (referring to the sampling location), was of particular interest to us, not only because it was not part of the training set for our tool, but also because of the scarcity of closely related genomes making this an outlier within the *Bacillota* phylum (**Figure 5D**). These challenges notwithstanding, our predictions for *Limnochorda* sp. EGM147 using COGs as features, selected for their biological interpretability and high predictive power, were coherent with enrichment culture growth conditions (**Table 2**). This shows that our models could be used to guide the selection of cultivation conditions for new isolates.

**Table 2.** Comparison between predicted and experimentally-defined environmental factor for Limnochorda sp. EGM147.

| Environmental factor | Observed | Predicted | Prediction method |
| --- | --- | --- | --- |
| Salinity | 3.5% | 3.8% | Regression |
| Temperature | 50.0°C | 47.7°C | Regression |
| Oxygen tolerance | Anaerobic | Anaerobic | Classification |
| pH | 7.3 | 7.1 | Regression |

### Molecular signatures of environmental adaptation

After developing ML models to predict salinity, temperature, oxygen, and pH, we explored the biological meaning of the important genomic features. We focused on the COGs and ncRNA families with the strongest association to the environmental parameters (**Figure 6**). Supporting our models and approach, several top features have relevant links in the literature, while others have indirect links that form promising candidates for future validation studies. These include non-coding RNA molecules (ncRNAs), molecules that are transcribed from the genome but are not translated into proteins, instead functioning directly as structural regulatory elements within the cell. In prokaryotes, ncRNAs play key roles in controlling gene expression, RNA processing and protein synthesis, often enabling rapid responses to environmental changes. Unlike protein-coding genes alone, ncRNAs can provide an additional, fast-acting layer of control that may be critical for fine-tuning cellular responses to environmental stresses such as salinity, temperature, oxygen, and pH.

**Figure 6.**
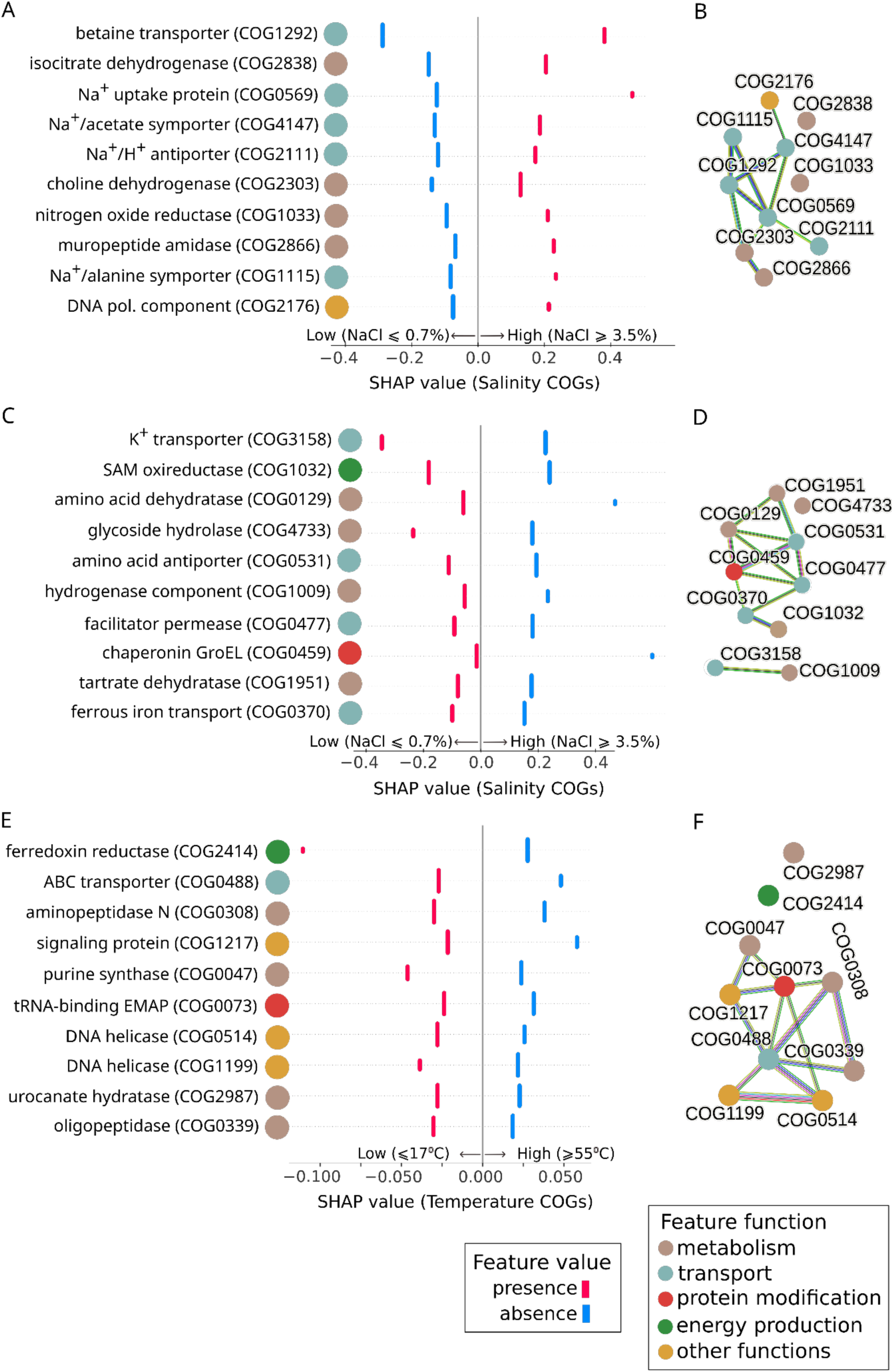
SHAP analysis and STRING connections of the most important COGs in ML classification models. **A, C, E**: beeswarm plots of local explanation for classification models of the top 10 most important COGs (from top to bottom) associated with high salinity **(AB)**, low salinity **(CD)**, and low temperature **(EF)**. COG colors indicate functional categories. **B, D, F**: STRING networks depicting known functional connections between COGs.

#### Temperature: heat-shock chaperones and translational delay

Several of the high ranking COGs in **Figure 6EF** have well-established temperature response mechanisms. This includes phenylalanyl-tRNA synthetase component COG0073, belonging to the small heat shock protein (HSP20) family, whose chaperone activity promotes thermotolerance in bacteria and archaea, protecting proteins from misfolding or aggregating under temperature stress (Li 2012). ATP-dependent DNA helicases COG0514 and COG1199 contribute to stabilizing the genome and repairing DNA damage in a temperature-dependent manner (Li 2010, Pantazaki 2008), and have a strong functional association (STRING combined score: 0.94, **Figure 6F**). Ribosome-associated GTPase COG1217 has chaperone activity. Mutations in the associated bipA gene induce cold sensitivity indicating it is cold-shock induced (Choi 2018).

Among ncRNAs, we identified relevant temperature-dependence of molecules including the Signal Recognition Particle (SRP). Three out of the four Rfam SRP families emerged as high-ranking features in the SHAP analysis of the temperature regression model: bacterial large SRP (Rfam RF01854, ranked 2nd), bacterial small SRP (RF00169, originally ranked 15th) and archaeal SRP (RF01857, 17th). We further investigated the association of these families with temperature by analyzing the conservation of their local genomic neighborhood. While the order *Bacteroidales* small SRP (RF04183, 43rd) ranked lower in the SHAP analysis, it was included to provide a comprehensive perspective on how the model interprets diverse structural variations of functionally similar small ncRNAs.

The SRP is part of a conserved ribonucleoprotein complex, important for the delay of translation and the translocation of proteins to the membrane (Zwieb 2005, Rosenblad 2009). SRPs may have diverse structures, with conservation of the so-called Helix 8 within the large SRP-specific (S) domain, which forms the binding site for the SRP54/Ffh protein (**Figure 7A**). This binding triggers a conformational change in the SRP complex, which signals the ribosome to slow down or pause protein synthesis. SRP variants are stratified by growth temperature (**Figure 7B**). For example, the archaeal and large bacterial SRPs are found in high-temperature isolates (87/87, 100% and 202/232, 87%, respectively), while the bacterial and *Bacteroidales* small SRPs are associated with low temperature (111/116, 96% and 294/359, 82%, respectively). This differentiation suggests a temperature adaptation related to translational delay. The bacterial large SRP typically includes the Alu domain, which is responsible for translational pausing (Halic 2004). In high-temperature environments, where the risk of co-translational misfolding is higher due to accelerated kinetics (Shalgi 2013), the ability of the large SRP to pause the ribosome may be a critical mechanism to allow for proper membrane protein insertion. This mechanism is consistent with the identification of chaperones among high ranking COGs.

**Figure 7.**
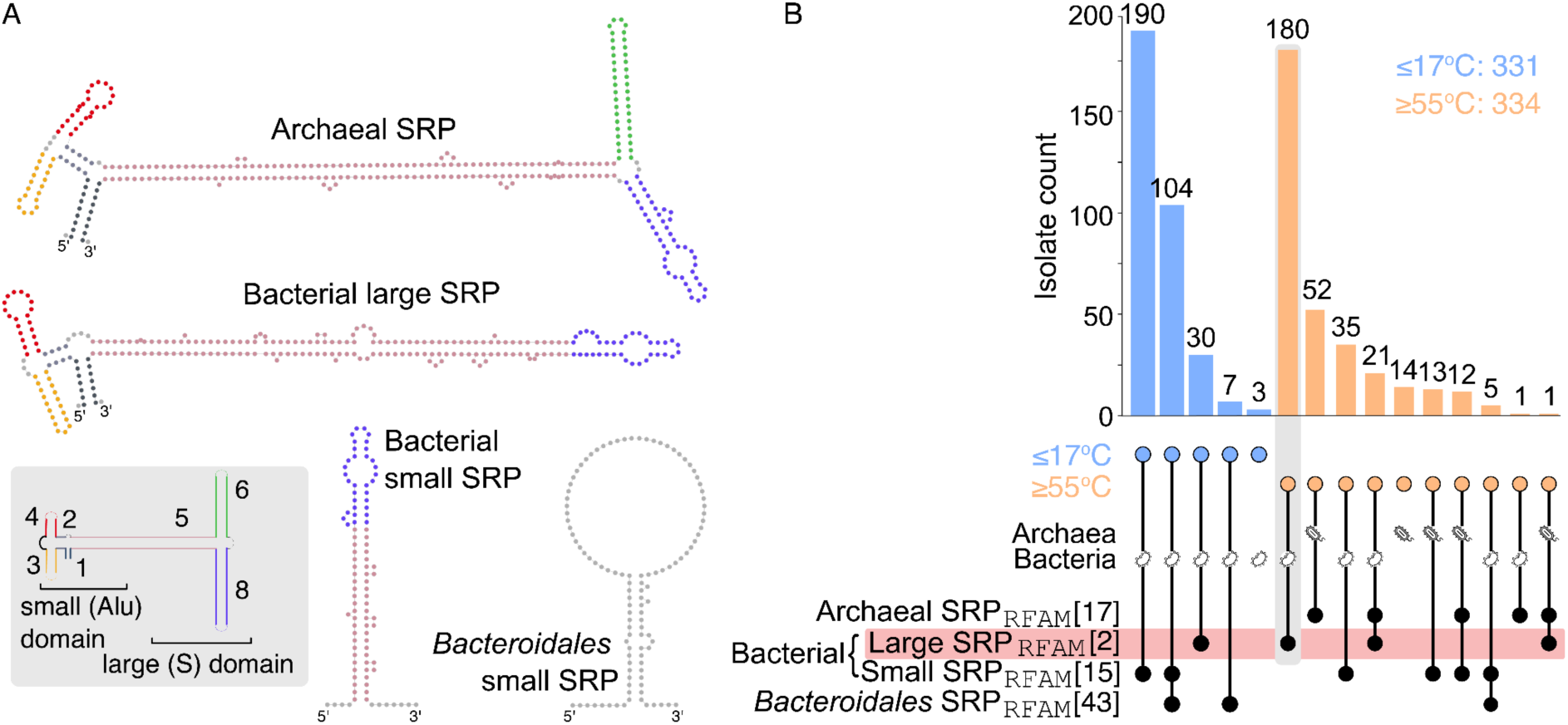
Structural diversity and temperature distribution of Signal Recognition Particle (SRP) RNA families. **(A)** Representative consensus secondary structures of the four SRP RNA variants. **(B)** Distribution of SRP variants across genomes in high and low temperature classes. The UpSet plot shows the presence and intersections of SRP RNA families in low- and high-temperature classes. The SHAP ranking of each family is shown in square brackets. Where the annotations of bacterial large and small SRPs overlapped on the same genome, the latter was removed.

#### Salinity: transport, metabolism, and hypoosmotic adaptation

In our salinity model, most of the important COGs are involved in transport and metabolism (**Figure 6A-D**). Known salinity adaptation mechanisms include a K⁺ transmembrane transporter (kup, COG3158), which promotes osmotic equilibrium in “salt-in” bacteria. In our model, COG3158 had negative SHAP values, indicating that its presence contributes to predicting the low-salinity class (**Figure 6C**). This corroborates previous observations that kup is more abundant in metagenomes from low-salinity environments (Wu 2024). Choline dehydrogenase (betA, COG2303), involved in the production of betaine, a key compatible solute that balances osmotic pressure in halophiles (Xia 2024, Yoo 2023), characterizes the high salinity class (**Figure 6A**). Dihydroxy-acid dehydratase (ilvD, COG0129) contributes to branched chain amino acid biosynthesis (Guez 2022), which promotes salinity-stress adaptation in plants (Sun 2024) and perhaps also in bacteria.

While much of the literature about salinity adaptation focuses on gene families such as ion transporters and osmolyte accumulation (Bremer and Krämer 2019, Zahra 2024, Chen 2017), our analysis also identified interesting ncRNAs. One specific antisense element that predicted salinity tolerance in low salt environments is anti-hemB (RF02837), which ranked within the top 20 most important ncRNA families in our SHAP analysis and was enriched in low-salinity isolates (**Figure 8A**). Our genomic context analysis showed that across proteobacterial families, anti-hemB is predominantly located upstream of the *hemB* gene (**Figure 8BC**), which it was proposed to regulate in cis (Weinberg 2007). The *hemB* gene encodes 5-aminolevulinic acid dehydratase, an enzyme that catalyzes the second step of heme biosynthesis and may therefore serve as a critical regulatory point for controlling the flux of this pathway (Franken 2011). Notably, anti-hemB is also in the proximity of cytochrome c biogenesis protein ResB (COG1333), which is essential for heme transport to apocytochromes (Jong-Hoon 2007). **Supplementary Figure 5** shows a comprehensive view of these genomic neighborhoods. Structural analysis of the ncRNA across this broad dataset reveals a highly conserved hairpin (**Figure 8D**), suggesting a critical functional role that is maintained across diverse taxa. This conservation in isolates where the ncRNA is decoupled from the *hemB* locus suggests that the functional motif may function in trans (Gottesman 2011), allowing the regulation of distal targets beyond its immediate genomic neighborhood. Our results support a role of anti-hemB in low salt conditions. Heme biosynthesis has previously been proposed to be reduced in high salinity (Sévin 2016), as bacteria may accumulate intermediates such as glutamate as osmoprotectant rather than consuming it for heme synthesis. In addition, lower heme biosynthesis may reduce iron demand, which could decrease ROS formation and limit oxidative damage under salt stress. Finally, heme biosynthesis may simply be costly, so downregulating it, for example with a regulatory ncRNA such as anti-hemB, could free up resources for osmoprotection (Sévin 2016, Aftab 2024).

**Figure 8.**
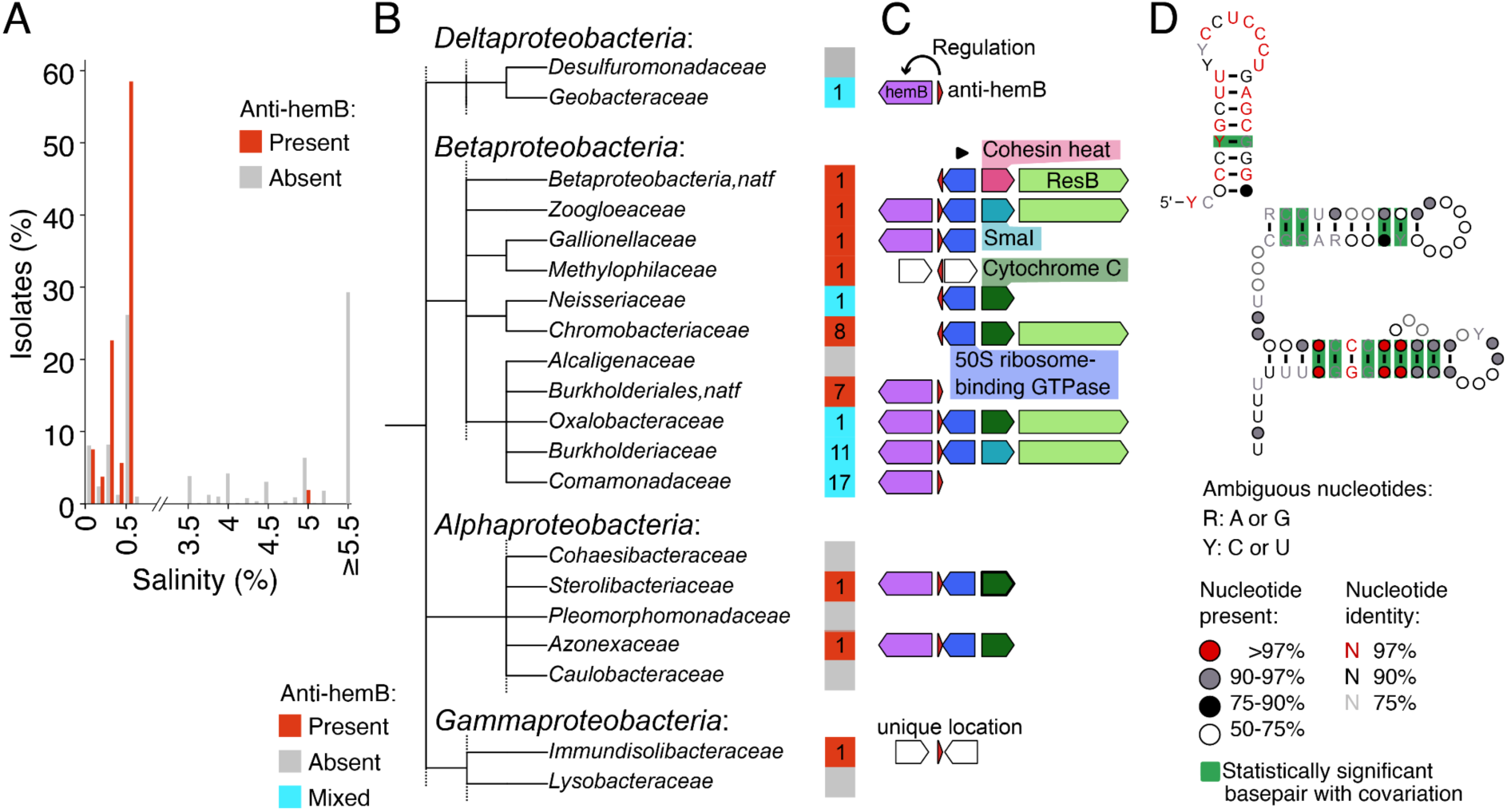
**(A)** Number of isolates with or without anti-hemB plotted against their salinity preferences. **(B)** Family-rank taxonomic tree with a color strip showing whether all or none of the strains in a family contain anti-hemB, or mixed. The number of anti-hemB containing strains is shown. **(C)** Synteny of the anti-hemB locus. The ncRNA (red triangle) is typically located upstream of hemB, which encodes 5-aminolevulinic acid dehydratase (see exceptions in **Supplementary Figure 5**). **(D)** Consensus secondary structure of anti-hemB across diverse taxa; colors indicate base-pair types and structural conservation.

#### Oxygen adaptation

Our classification model for aerobic versus anaerobic isolates prioritised interesting COGs. First, the FAD-linked oxidase domain glycolate oxidase COG0277, involved in redox balance and oxidative metabolism (Chen 2021). Second, glutamate dehydrogenase COG2902, involved in catabolising glutamate to 2-oxoglutarate, could have an indirect role in oxygen mechanisms, considering that an enzyme dependent on 2-oxoglutarate has been identified as an oxygen sensor in a recent hypoxia study (Lee 2025). Third, cytochrome c oxidase subunit COG0843, with heme-copper oxidase activity, a key component of the electron transport chain in aerobic respiration (Centeno-Leija 2013, de Gier 1994), is associated with aerobic isolates in our model (**Supplementary Figure 6AB**). Fourth, we found the radical SAM superfamily enzyme COG1032, which was also important for salinity classification. SAM is a co-substrate for different enzymatic reactions including radical mediated processes, and SAM enzymes occur in anaerobic bacteria (Benjdia 2017) and archaea in oxygen-deficient environments (Li 2024). They catalyze radical chemistry and are oxygen-sensitive (Vey and Drennan 2018). Our SHAP analysis corroborates these findings, with the presence of COG1032, being associated with the anaerobic phenotype (**Supplementary Figure 6CD**).

#### pH adaptation

Important COGs in our pH regression model are related to transport, metabolism, and protein modification (**Supplementary Figure 6EF**). The ATP-dependent Clp protease ATP-binding subunit COG0542 is part of a pH-dependent ATP-dependent protease complex (Kim 2022). The simple sugar transporter ATP-binding protein COG1129 belongs to the ABC transporter superfamily involved in sugar transport, potentially contributing to solute homeostasis in the cell (Jeckelmann and Erni 2020). ABC-2 type drug efflux transport system protein COG0842 is a permease component involved in stress responses and extrusion mechanisms (Zaide 2008). COG2208 is a metal-dependent phosphoserine phosphatase involved in sensing and responding to environmental changes by performing protein conformational changes that affect ligand interaction (Treuner-Lange 2010). Finally, molybdopterin containing dehydrogenase subunit COG4631 functions e.g. in xanthine dehydrogenase, which is involved in the uric acid production with the reduction of NAD^+^ to NADH (Wang 2016).

## Discussion

We developed the FxTractor pipeline to automatically extract diverse features encoded on prokaryotic genome sequences (**Figure 2**). FxTractor is open-source, easy to use, and built on the extendable Snakemake framework that allows users to readily implement additional features of interest. A tutorial covering installation and use, with example files, is available at https://github.com/MGXlab/FxTractor, where we invite users to contribute by adding additional features to the open source code through pull requests, raising issues, or requesting new features.

We used FxTractor to extract different feature types from 13,554 isolates, including eggNOG clusters of orthologous groups (COGs), ncRNA families, nucleotide usage (9-nucleotide kmers), and amino acid frequencies. We then built ML models to predict environmental factors (**Figure 1A**) and validated them with held-out testing data, as well as the newly sequenced genome of the extremophile *Limnochorda* sp. EGM147 (**Figure 5**). We obtained predicted values that closely aligned with the conditions used during laboratory cultivation (**Table 2**), suggesting that FxTractor models could be used to guide the cultivation of new isolates.

Next, we were interested in comparing different feature types in terms of their predictive power. The literature shows that different feature types can be used to make predictions of preferred growth conditions. Well-performing models based on amino acid frequencies or amino acid descriptors have been developed (Colette 2025, Li 2019, Barnum 2024). Successful attempts were also made using functional features such as protein sequences (Gado 2025), Pfam annotations (Koblitz 2025, Weimann 2016, Ramoneda 2023), COGs (Alneberg 2020, Wu 2024), KEGG orthologs (Barberán 2017), or oxygen utilizing enzymes (Jabłońska 2019). In a direct comparison, amino acid frequencies outperformed protein isoelectric point frequencies (Barnum 2024). Another work showed that Pfam and eggNOG-based annotations performed better than other types of functional annotation (Koblitz 2025). NcRNA families have not yet been explored, with only tRNA and rRNA being used as features to improve models based on genomic sequences for temperature prediction (Sauer 2019).

Our classification models showed similar performance for all feature types, although ncRNAs performed slightly less well than the other feature types (**Figure 4A-C**). This could partly be due to the lower sequence-level conservation of ncRNAs, which function as folded RNA structures, making them more difficult to detect. Another possible explanation for the poorer performance of ncRNAs could be the lower number of included features (n=1,110, see **Table 1**) relative to COGs and kmers, giving them less resolution for the predictions. However, we note that the 20 amino acid frequencies rather had slightly higher prediction accuracies in our regression models than other feature types, while kmers (n=131,072) performed relatively less well (**Figure 4D-F**). Amino acid frequencies thus seem to respond strongly to temperature and salinity. Indeed, their frequencies have been considered hallmark signatures for high-salt-adapted organisms, with the enriched negatively charged residues thought to improve the ability of proteins to remain soluble and avoid aggregation (Shen 2024). Still, annotation-based features, such as COGs or ncRNA families, offer more direct insight into underlying biological mechanisms.

By including additional feature types and predicting diverse phenotypes or environmental parameters, we expect that future research leveraging the FxTractor pipeline for hypothesis testing will identify novel molecular mechanisms, including environmental adaptation. Our highlighted examples demonstrate that important protein and ncRNA families may be readily revealed, thus improving the understanding of the relationship between microbes and their surrounding environment.

## Acknowledgements

We thank Peter Stadler and Zasha Weinberg for valuable discussions on ncRNAs.

## Author contributions

**MBWC**: conceptualization, software, data curation, writing - original draft, visualization. **RB**: conceptualization, software, writing - review & editing. **MS**: software, writing - original draft, visualization. **AL**: software, writing - review & editing. **FB**: investigation, writing - review & editing, visualization. **HF**: software, visualization, funding acquisition. **CRJH**: resources, funding acquisition, writing - review & editing. **MM**: resources, writing - review & editing, funding acquisition. **BED**: resources, conceptualization, writing - original draft, writing - review & editing, supervision, funding acquisition.

## Data availability

The data that support the findings of this study can be found in https://github.com/MGXlab/abiotic_environment. The FxTractor pipeline is available at https://github.com/MGXlab/FxTractor.

## Supplementary Materials

### Supplementary Figures

**Supplementary Figure 1.**
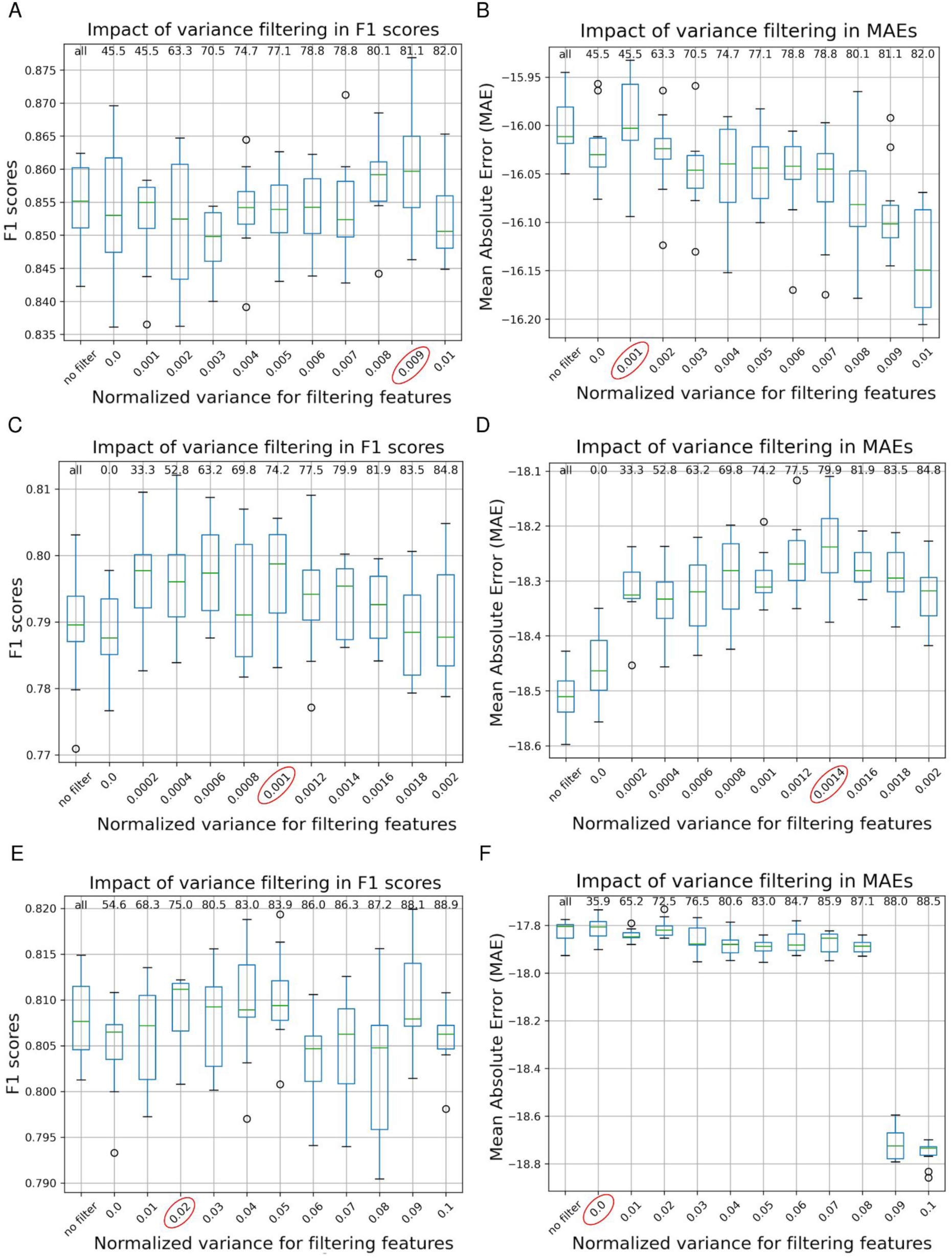
Histograms of RF performances for different thresholds of variance. Datasets for salinity are shown as examples. **AB**: COGs, **CD**: kmers, **EF**: ncRNAs. Numbers on the top of the plot indicate the percentage of features filtered out using the threshold (“all” indicates all features were kept). Red circles indicate the chosen threshold. **ACE**: classification models, **BDF**: regression models.

**Supplementary Figure 2.**
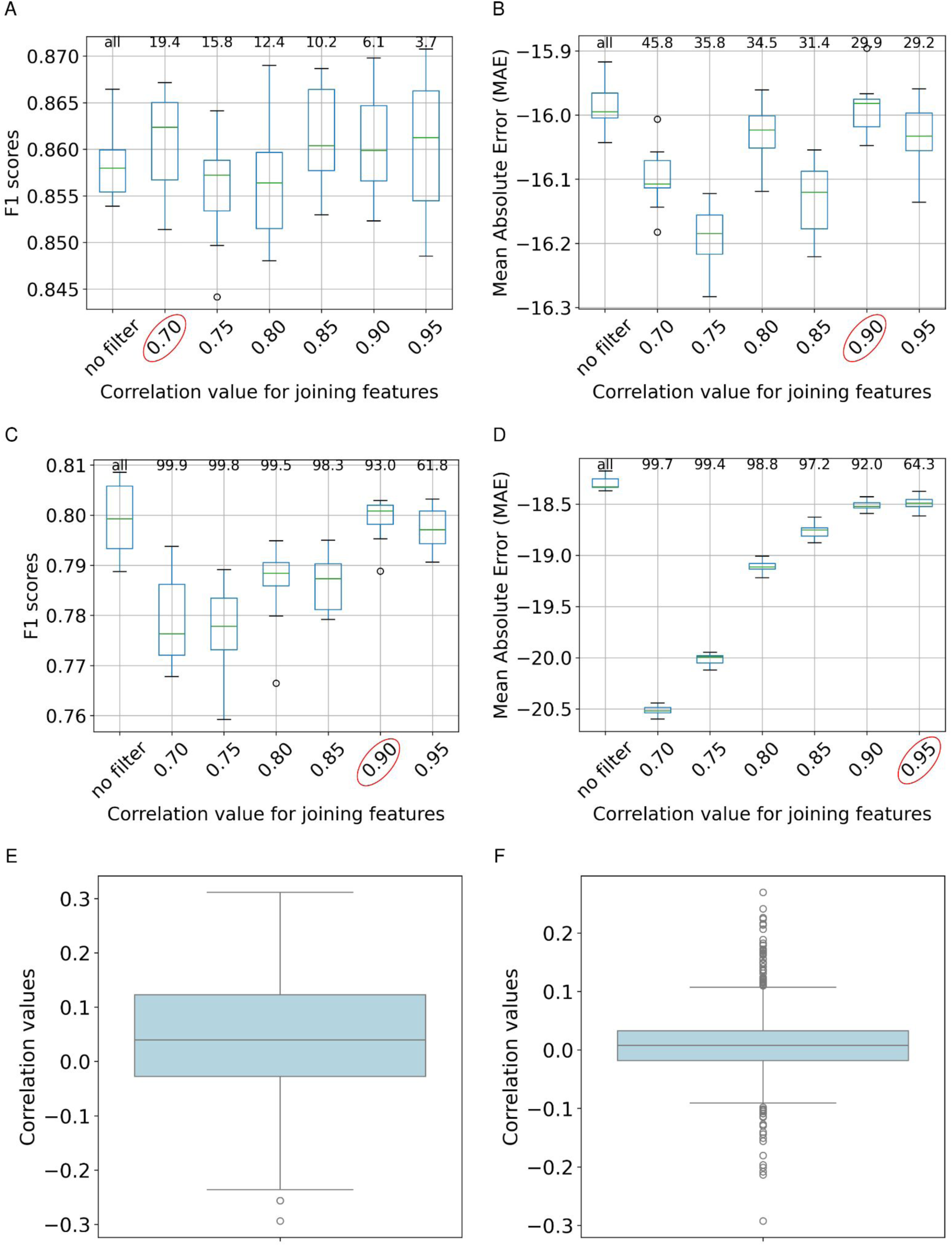
Results of Spearman correlation analysis for different types of features. Datasets for salinity are shown as examples. **A-D**: Histograms of RF performances for different thresholds of Spearman correlation. **AB**: COGs, **CD**: kmers. **A-D**: numbers on the top of the plot indicate the percentage of features filtered out using the threshold (“all” indicates all features were kept). **AC**: classification models, **BD**: regression models. **EF**: For ncRNA families, no features were clustered due to the low correlation between features. **E**: classification model, **F**: regression model.

**Supplementary Figure 3.**
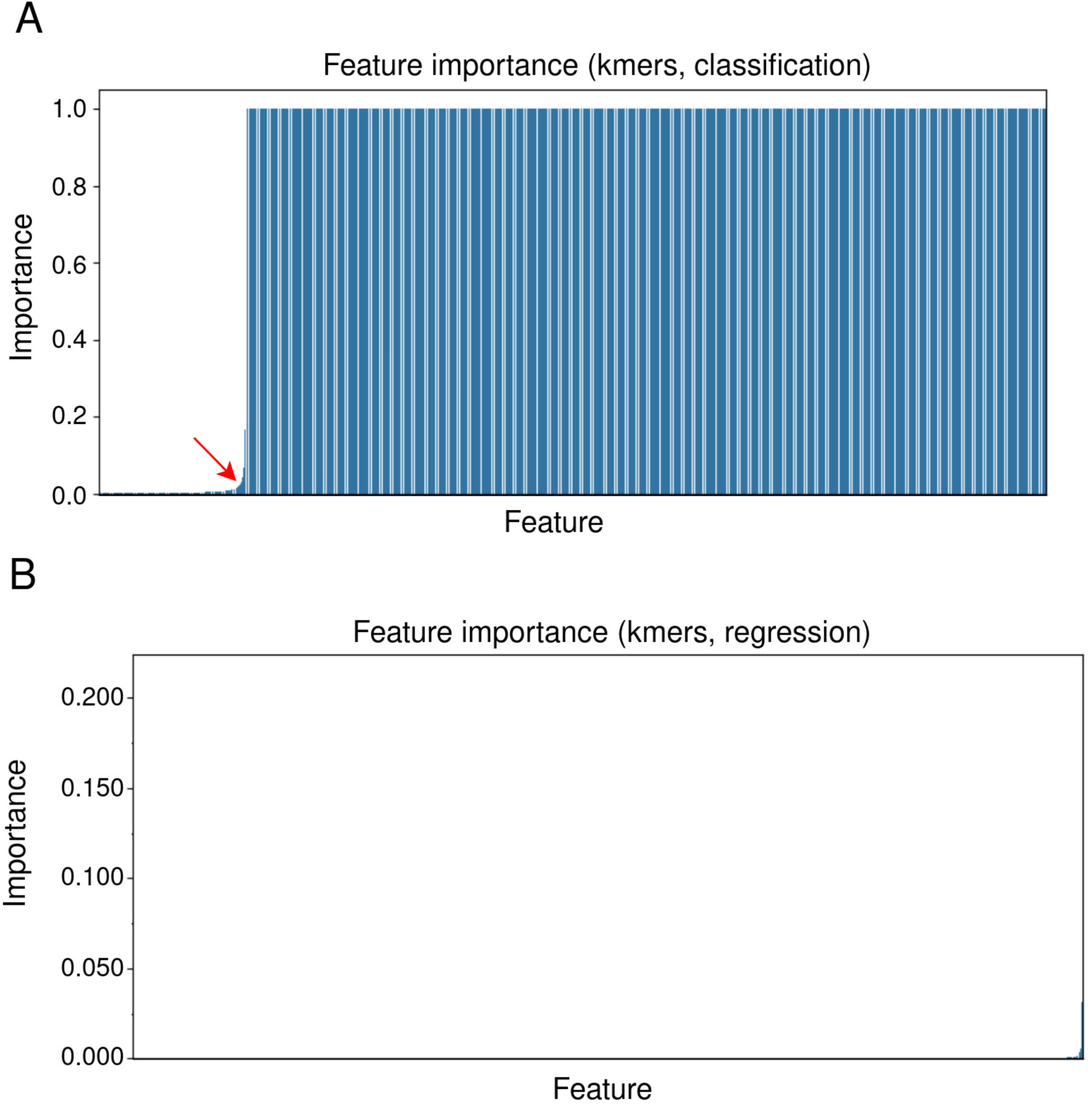
Results of datasets analysed with RFE, with temperature given as example. **A**: classification model of kmers, with “elbow point” marked with a red arrow, **B**: regression models of kmers, with no visible “elbow point”. X-axis: features ranked by importance scores, y-axis: importance scores calculated with decision trees.

**Supplementary Figure 4.**
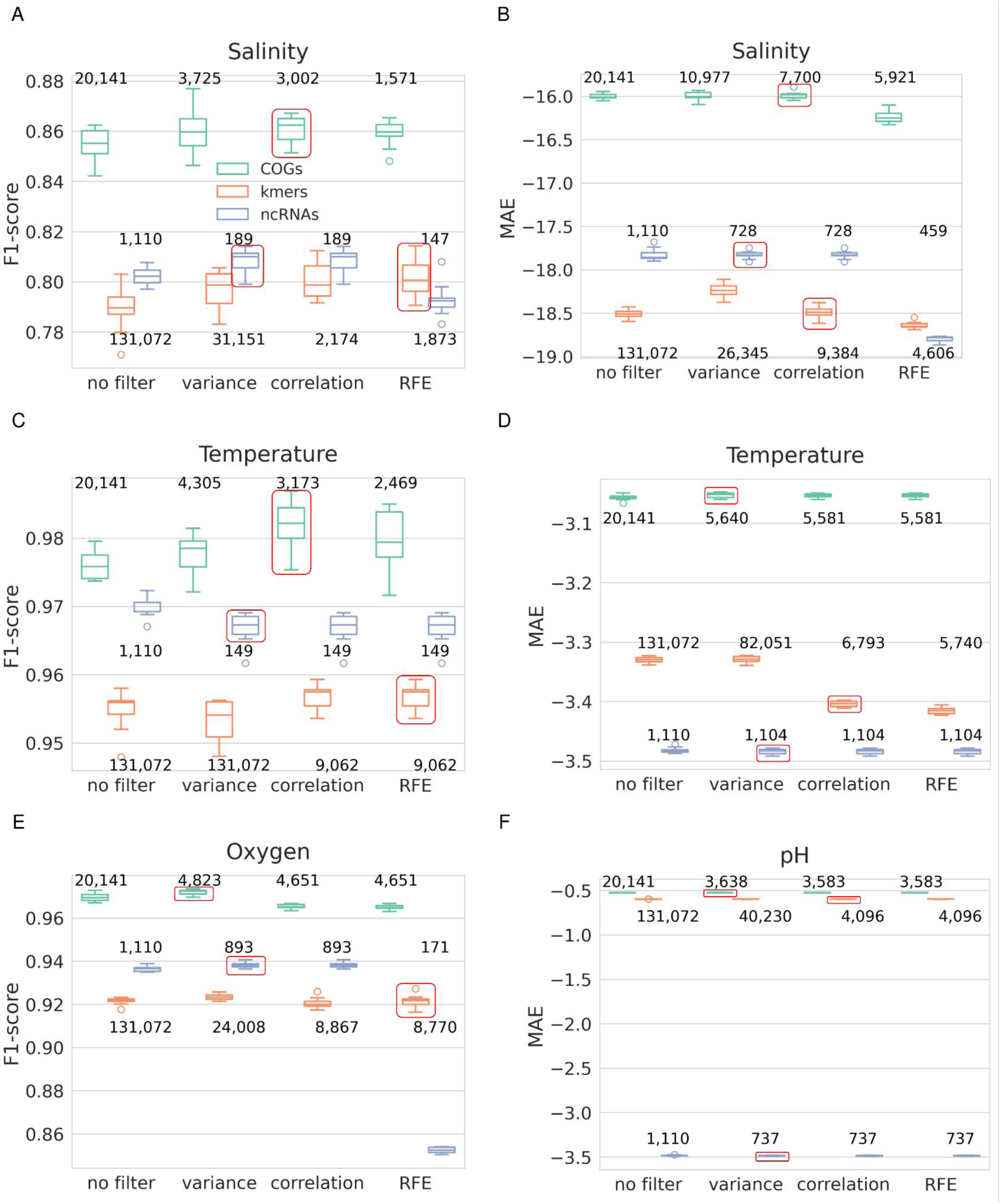
Performances of Random Forest (RF) models produced with datasets of each selection step. **AB**: salinity, **CD**: temperature, **E**: oxygen, **F**: pH, **ACE**: classification models, **BDF**: regression models. Feature types are shown in different colors, with green: gene families (COGs), blue: ncRNA families and orange: kmers (as specified in panel **A**). The final chosen datasets are marked with red squares in the plots.

**Supplementary Figure 5.**
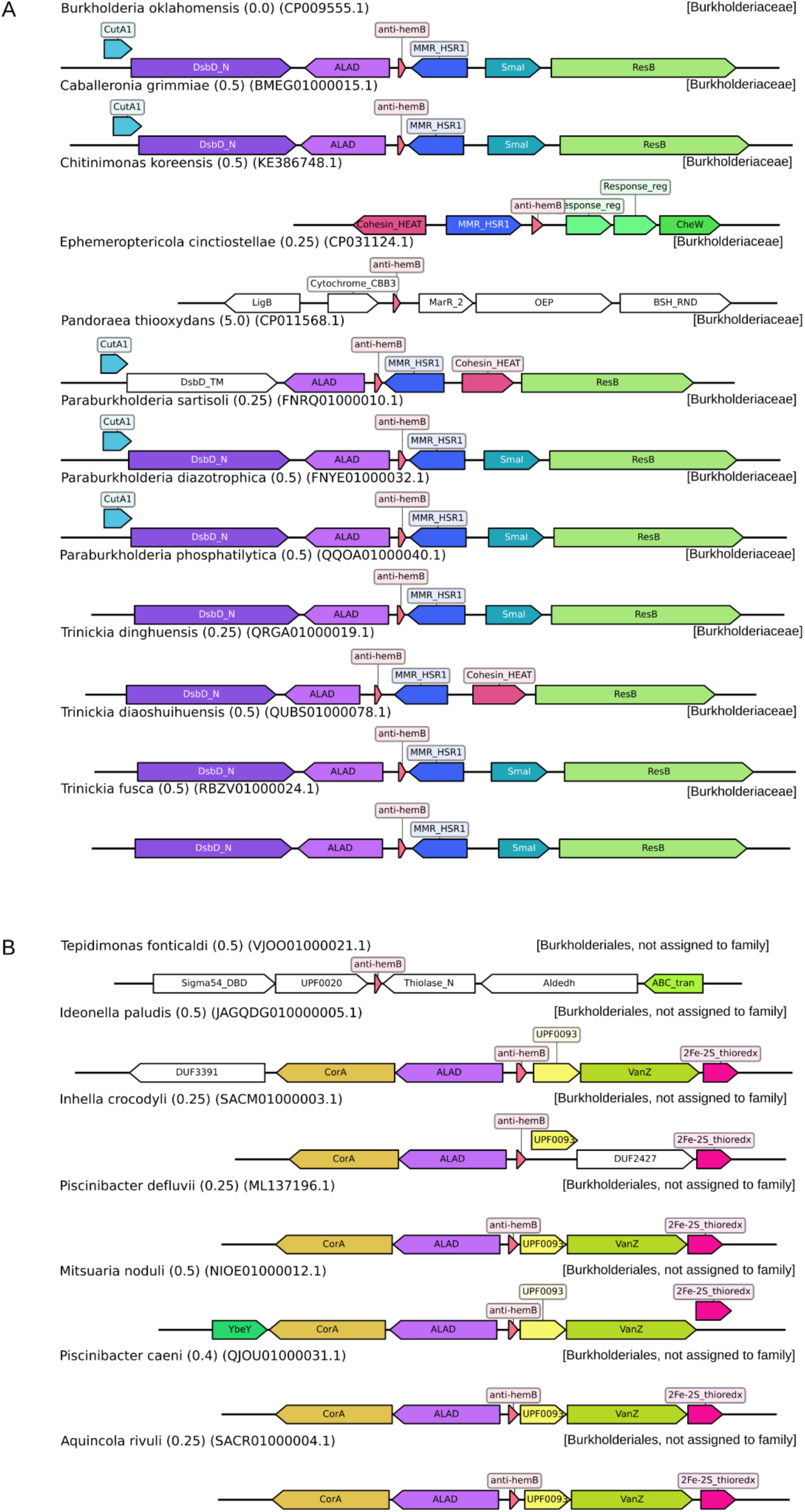
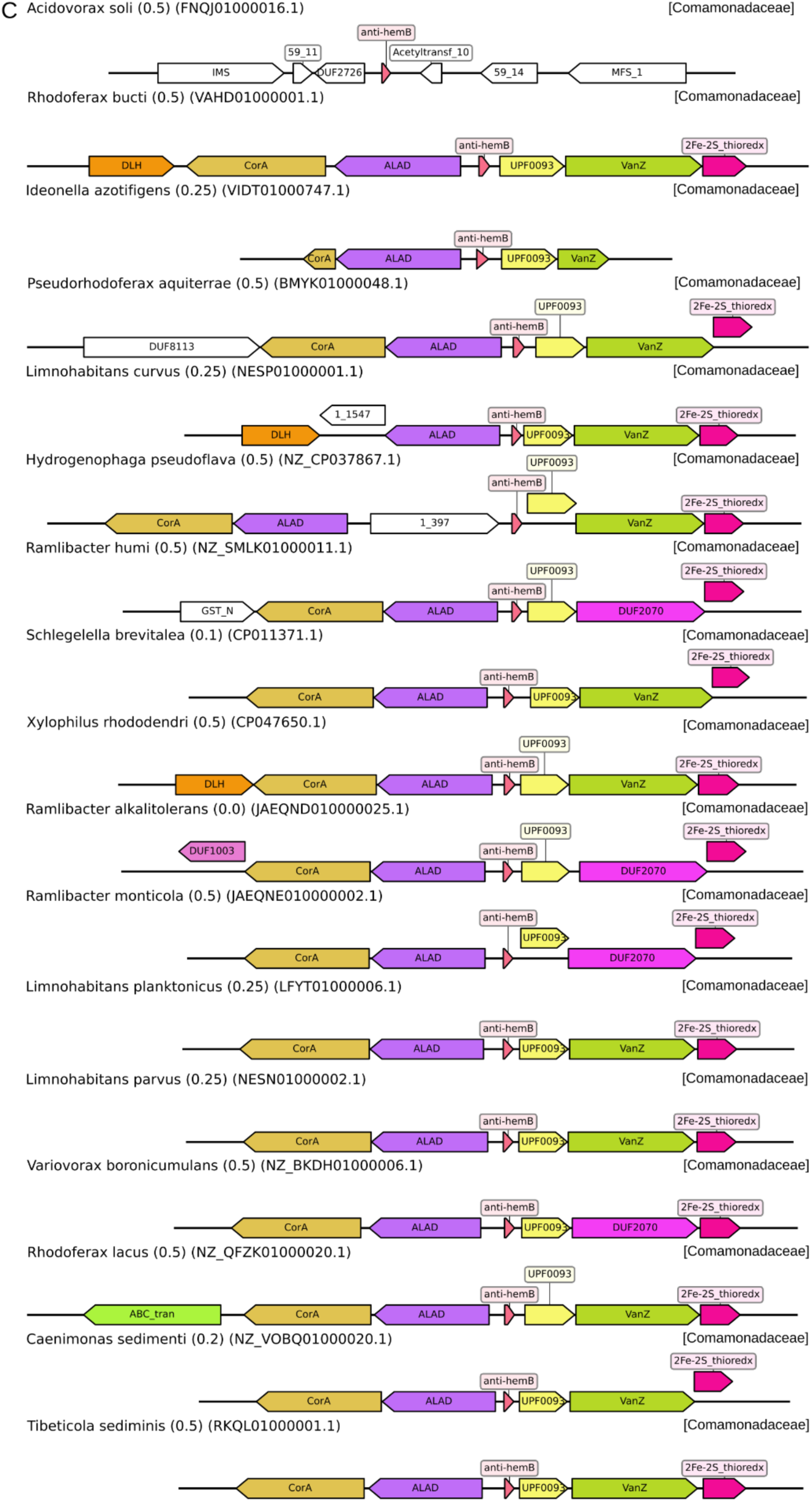
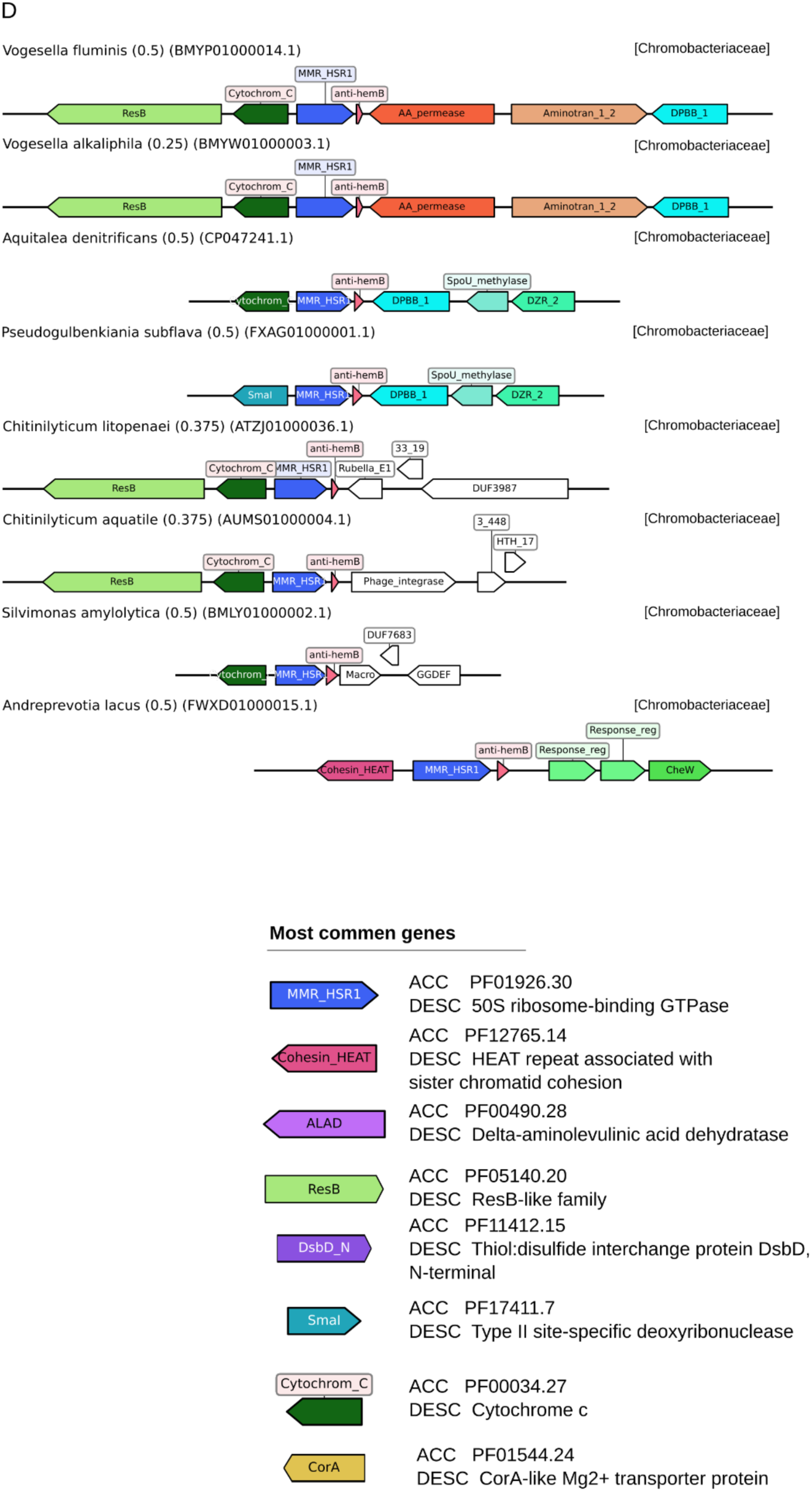
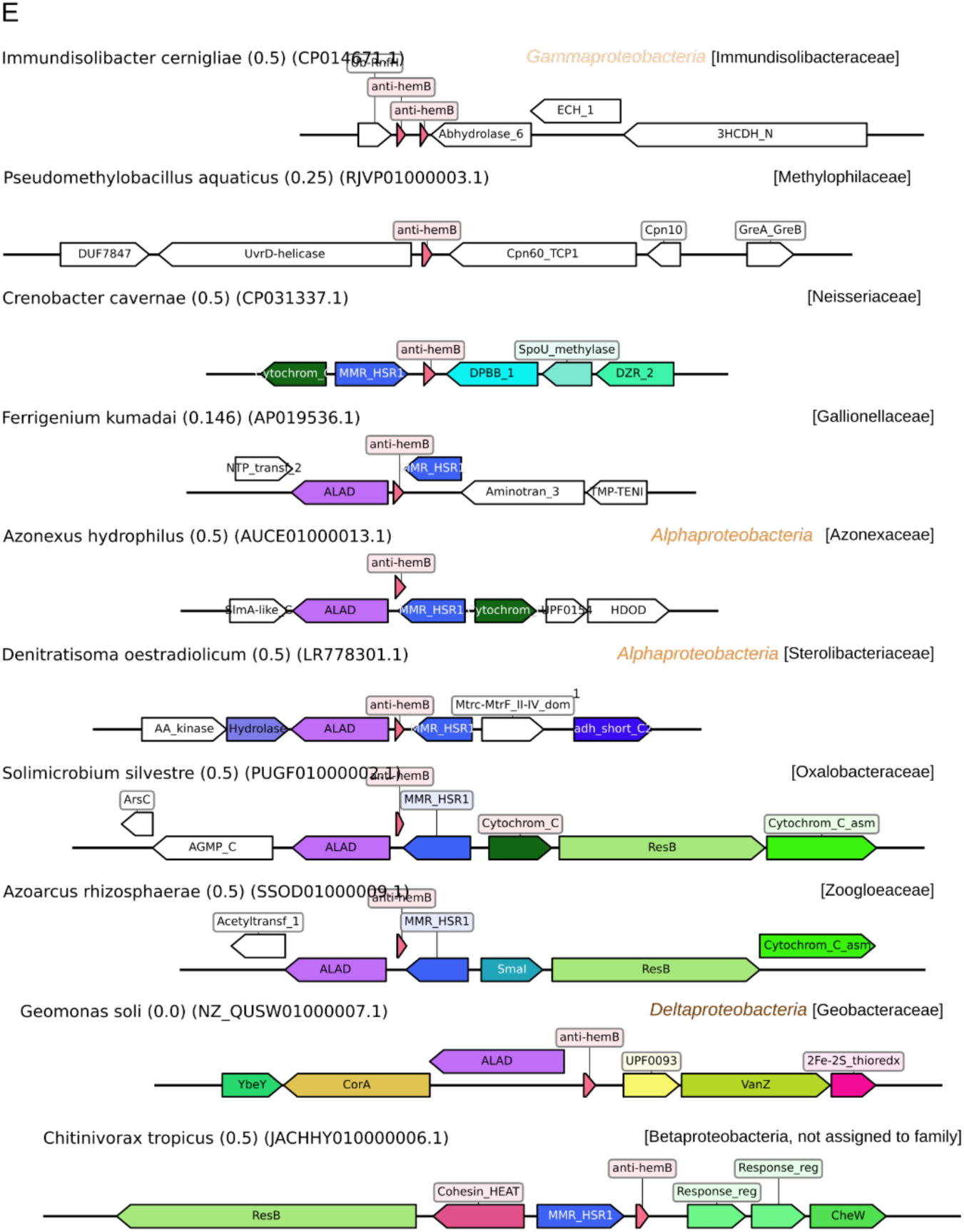
Comparative analysis of the genomic neighborhoods surrounding the anti-hemB ncRNA. Representative genomic loci from multiple bacterial assemblies are shown, centered on the anti-hemB regulatory RNA (small red arrows). Genes are represented by colored arrows indicating relative position and orientation. A high degree of syntenic conservation is observed across taxa, with **A**: Burkholderiaceae, **B**: Burkholderiales, not assigned to family, **C**: Commanadaceae, **D**: Chromobacteriaceae, **E**: others. Divergent flanking regions are indicated by white colored arrows, representing less conserved unique genomic context. The most common and important gene family assignments are provided in the legend in part D (bottom right).

**Supplementary Figure 6.**
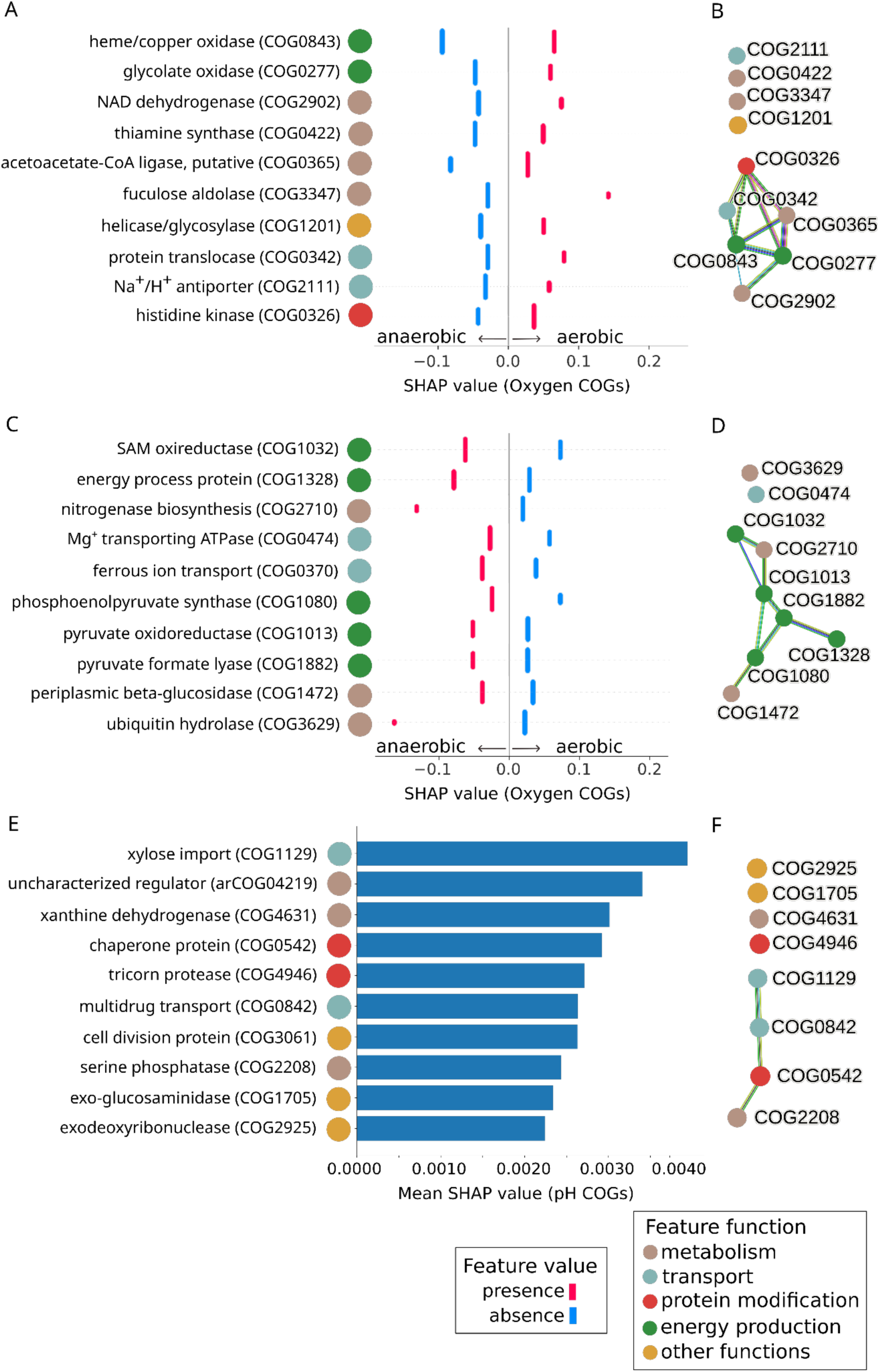
SHAP analysis and STRING connections of the most important COGs in ML models. **A, C**: beeswarm plots of local explanation for classification models for the top 10 most important COGs (from top to bottom) associated with aerobic (**AB**) and anaerobic (**CD**) conditions. **E:** mean SHAP values of the top 10 (from top to bottom) COGs of the regression model for pH (**EF**). COG colors indicate functional categories. **B, D, F**: STRING networks depicting known functional connections between COGs.

### Supplementary Tables

***Supplementary Table 1.** List of all 91,228 microbial isolates (90,241 bacteria and 987 archaea) whose genomes were downloaded from BacDive (Schober 2025), including metadata fields (GC content, salinity, temperature, pH, oxygen tolerance) and full taxonomic lineage. The last column indicates in which ML models the genome was used.*

**Supplementary Table 2.**
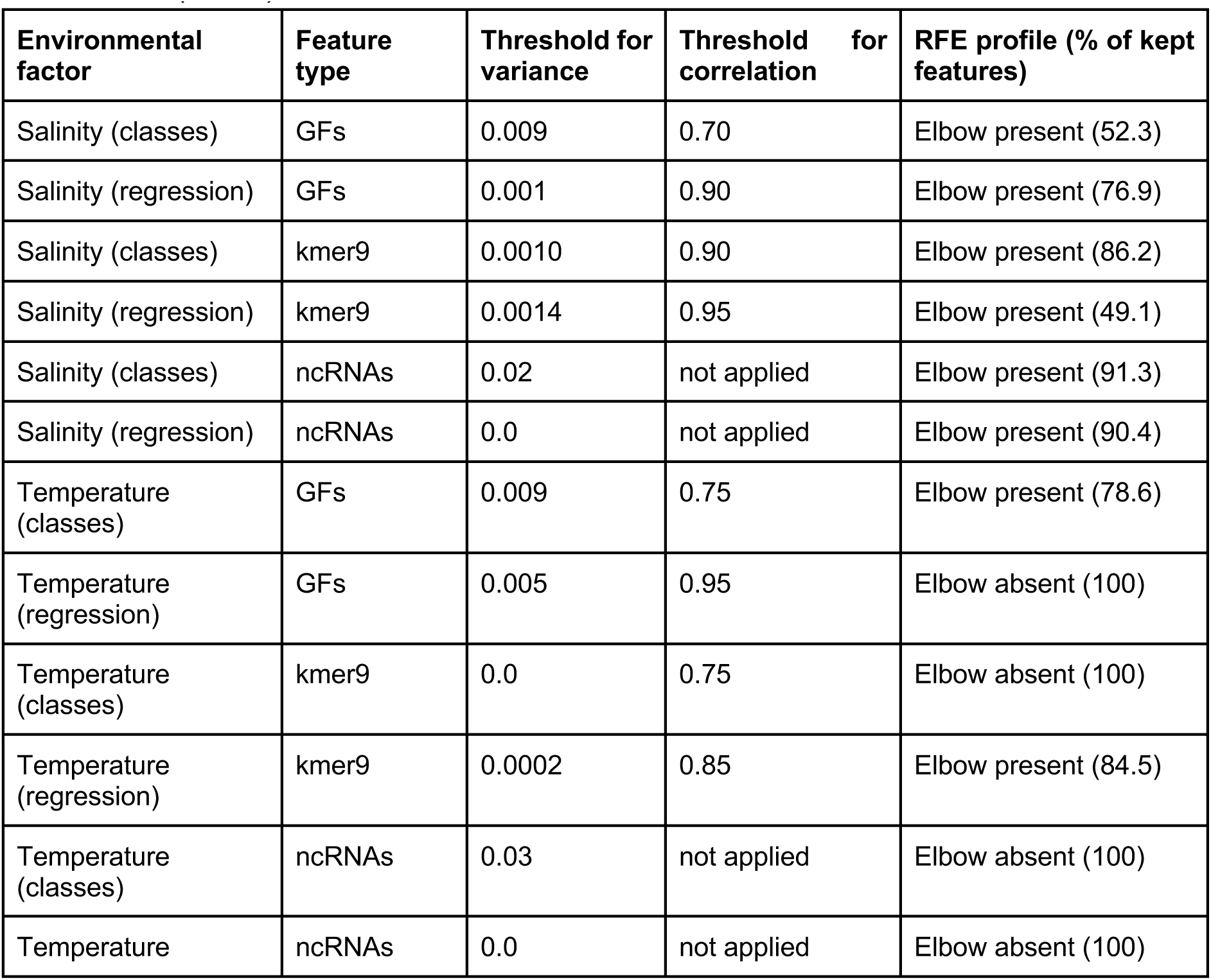
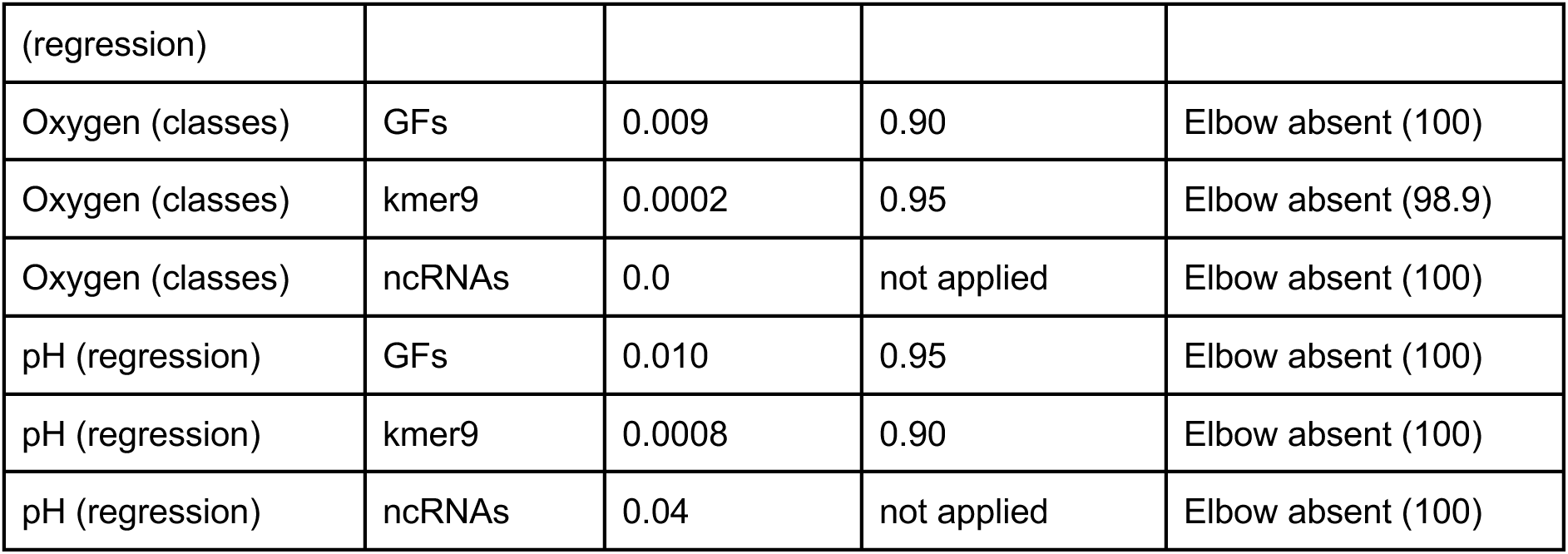
Feature selection summary. Appropriate thresholds were chosen for variance and correlation steps considering feature type and ML classification or regression. For the third step of selection, RFE, the plot profile is given along with the percent of features kept (taking into account the previous selection step: correlation). No feature was clustered by correlation for ncRNAs, since all had low correlations (<0.60).

**Supplementary Table 3.** Number of features selected per selection step and dataset. Text colored in blue indicates the initial number of features, while text in red indicates the final number of features. The number of isolates in classification datasets combines the two contrasting classes (low and high factor value). The final datasets were chosen as the ones with the highest performance in benchmarking different selection steps (**Supplementary Figure 4**). Exceptions to this rule are marked with an asterisk: files for kmer regression.

| Environmental factor | Feature type | Model type | Isolates | Features | Variance | Correlation | RFE |
| --- | --- | --- | --- | --- | --- | --- | --- |
| Salinity | COGs | Clas. | n = 932 | 20,141 | 3,725 | 3,002 | 1,571 |
|  | COGs | Reg. | n = 3,418 | 20,141 | 10,977 | 7,700 | 5,921 |
|  | kmers | Clas. | n = 932 | 131,072 | 31,151 | 2,174 | 1,873 |
|  | kmers | Reg. | n = 3,418 | 131,072 | 26,345 | 9,384* | 4,606 |
|  | ncRNAs | Clas. | n = 932 | 1,110 | 189 | 189 | 147 |
|  | ncRNAs | Reg. | n = 3,418 | 1,110 | 728 | 728 | 459 |
| Temperature | COGs | Clas. | n = 668 | 20,141 | 4,305 | 3,173 | 2,469 |
|  | COGs | Reg. | n = 13,198 | 20,141 | 5,640 | 5,581 | 5,581 |
|  | kmers | Clas. | n = 668 | 131,072 | 131,072 | 9,062 | 9,062 |
|  | kmers | Reg. | n = 13,198 | 131,072 | 82,051 | 6,793* | 5,740 |
|  | ncRNAs | Clas. | n = 668 | 1,110 | 149 | 149 | 149 |
|  | ncRNAs | Reg. | n = 13,198 | 1,110 | 1,105 | 1,105 | 1,105 |
| Oxygen | COGs | Clas. | n = 3,402 | 20,141 | 4,823 | 4,651 | 4,651 |
|  | kmers | Clas. | n = 3,402 | 131,072 | 24,008 | 8,867 | 8,770* |
|  | ncRNAs | Clas. | n = 3,402 | 1,110 | 893 | 893 | 171 |
| pH | COGs | Reg. | n = 3,630 | 20,141 | 3,638 | 3,583 | 3,583 |
|  | kmers | Reg. | n = 3,630 | 131,072 | 40,230 | 4,096* | 4,096 |
|  | ncRNAs | Reg. | n = 3,630 | 1,110 | 737 | 737 | 737 |

**Supplementary Table 4.**
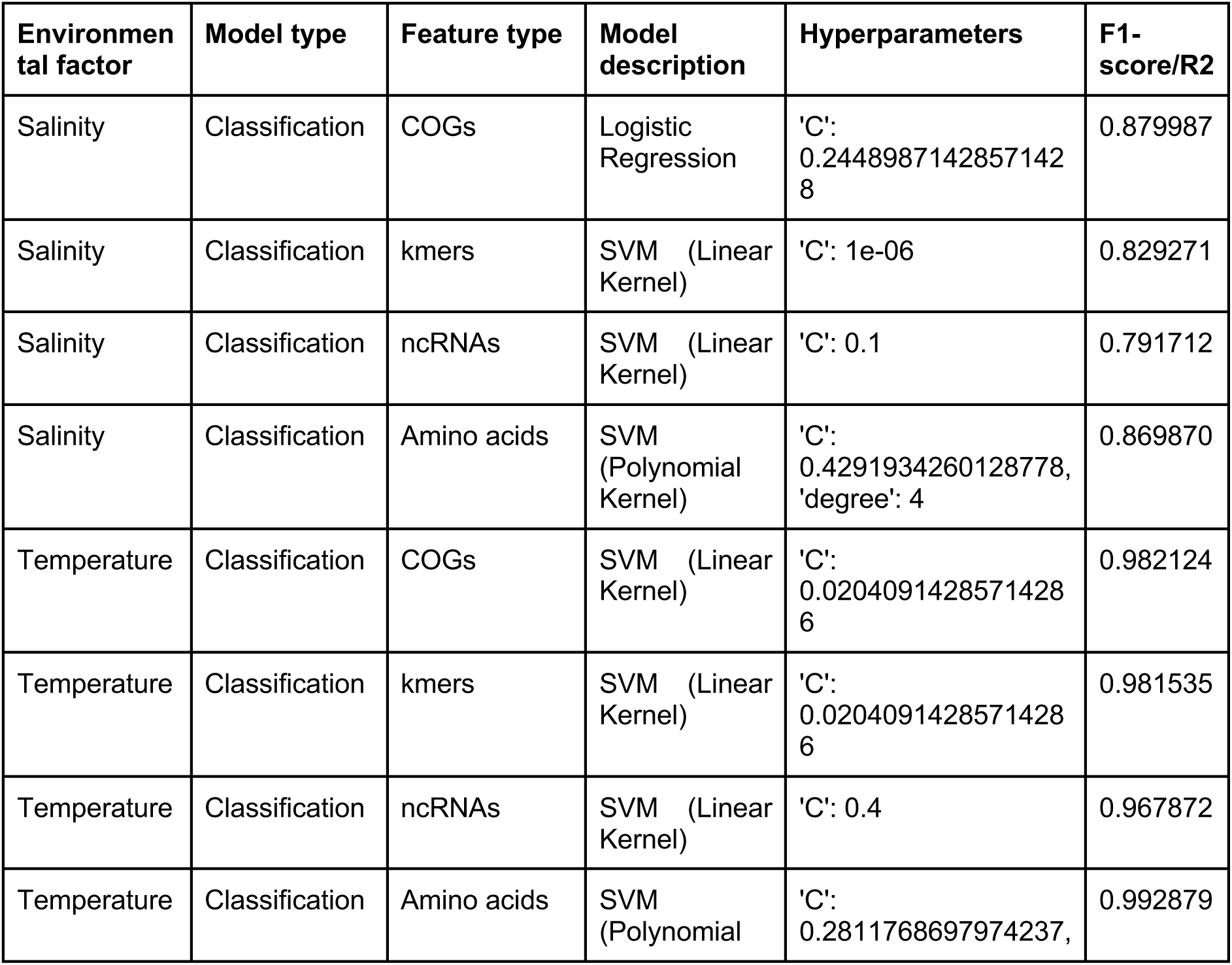
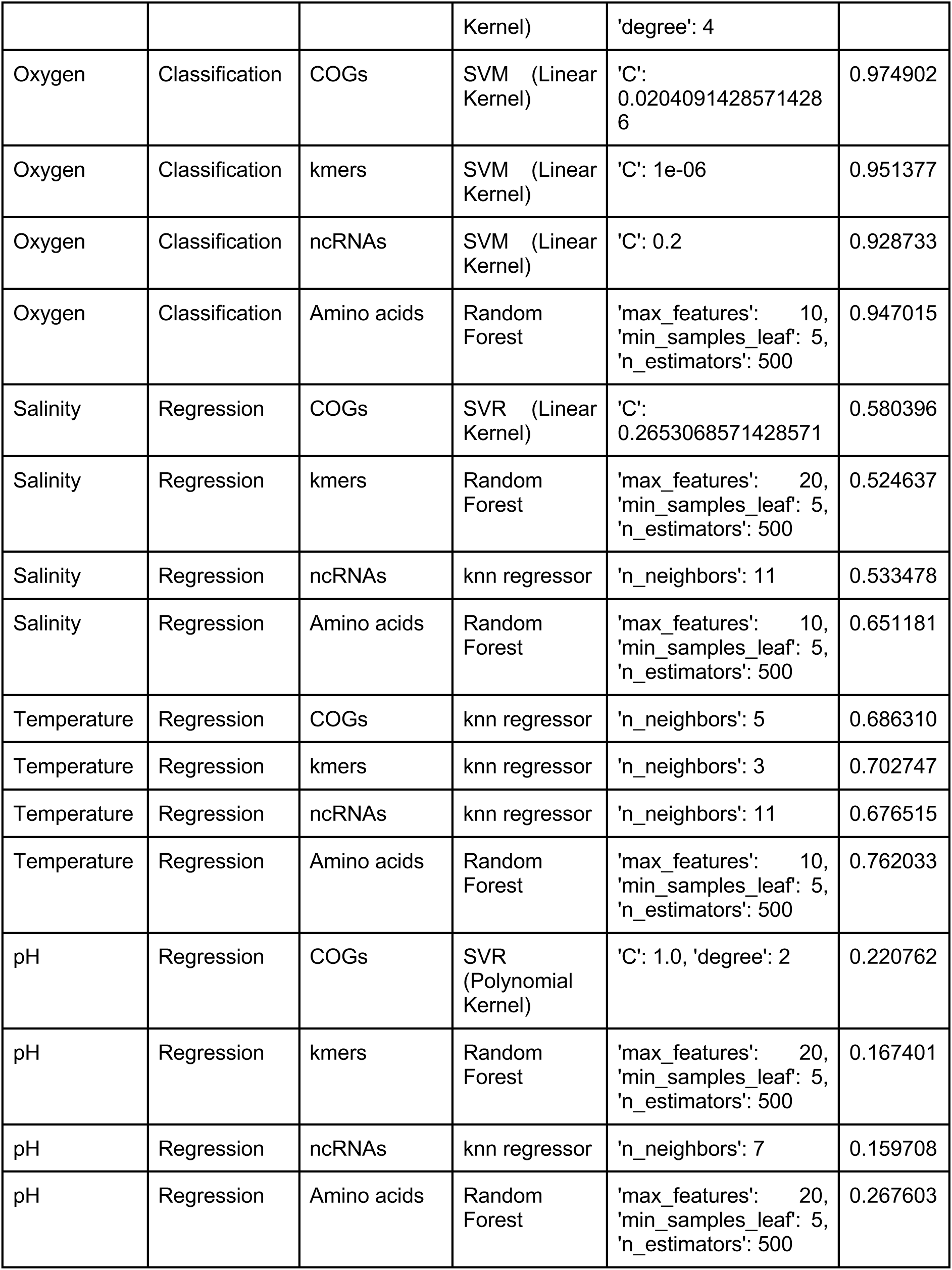
Best models per dataset (considering environmental factor, feature type and classification or regression) after hyperparameter tuning.

### Supplementary Files

Rankings of SHAP analysis and alignment files of ncRNAs can be accessed at: https://github.com/MGXlab/abiotic_environment.

